# scMeFormer: a transformer-based deep learning model for imputing DNA methylation states in single cells enhances the detection of epigenetic alterations in schizophrenia

**DOI:** 10.1101/2024.01.25.577200

**Authors:** Jiyun Zhou, Chongyuan Luo, Hanqing Liu, Matthew G. Heffel, Richard E. Straub, Joel E. Kleinman, Thomas M. Hyde, Joseph R. Ecker, Daniel R. Weinberger, Shizhong Han

**Affiliations:** Lieber Institute for Brain Development, Johns Hopkins Medical Campus, Baltimore, MD, 21287, USA; Department of Human Genetics, University of California, Los Angeles, Los Angeles, CA, 90095, USA; Genomic Analysis Laboratory, The Salk Institute for Biological Studies, La Jolla, CA, 92037, USA; Bioinformatics Interdepartmental Program, University of California Los Angeles, Los Angeles, CA 90095, USA; Howard Hughes Medical Institute, The Salk Institute for Biological Studies, La Jolla, CA, 92037, USA; Department of Psychiatry and Behavioral Sciences, Johns Hopkins University School of Medicine, Baltimore, MD, 21287, USA; Department of Genetic Medicine, Johns Hopkins University School of Medicine, Baltimore, MD 21205, USA; Department of Neuroscience, Johns Hopkins University School of Medicine, Baltimore, MD, 21205, USA; Department of Neurology, Johns Hopkins University School of Medicine, Baltimore, MD, 21205, USA

## Abstract

DNA methylation (DNAm), a crucial epigenetic mark, plays a key role in gene regulation, mammalian development, and various human diseases. Single-cell technologies enable the profiling of DNAm states at cytosines within the DNA sequence of individual cells, but they often suffer from limited coverage of CpG sites. In this study, we introduce scMeFormer, a transformer-based deep learning model designed to impute DNAm states for each CpG site in single cells. Through comprehensive evaluations, we demonstrate the superior performance of scMeFormer compared to alternative models across four single-nucleus DNAm datasets generated by distinct technologies. Remarkably, scMeFormer exhibits high-fidelity imputation, even when dealing with significantly reduced coverage, as low as 10% of the original CpG sites. Furthermore, we applied scMeFormer to a single-nucleus DNAm dataset generated from the prefrontal cortex of four schizophrenia patients and four neurotypical controls. This enabled the identification of thousands of differentially methylated regions associated with schizophrenia that would have remained undetectable without imputation and added granularity to our understanding of epigenetic alterations in schizophrenia within specific cell types. Our study highlights the power of deep learning in imputing DNAm states in single cells, and we expect scMeFormer to be a valuable tool for single-cell DNAm studies.

## Introduction

DNA methylation (DNAm), a fundamental epigenetic mechanism involving the addition of a methyl group to cytosines, plays a crucial role in gene regulation, mammalian development, and various human diseases^1^. Single-cell technologies enable the profiling of DNAm states at cytosines within the DNA sequence of individual cells, which complements single-cell transcriptome studies in understanding cellular heterogeneity, developmental processes, and disease states^2, 3^. However, due to the limited DNA material available from individual cells and inherent technical limitations, current technologies often suffer from sparse coverage for CpG sites, typically measuring less than 10% of CpG sites in a single cell^4–7^. This limits their full potential to uncover the epigenetic landscape at single-cell resolution.

To address the challenge of sparse coverage in the single-cell DNAm dataset, computational methods have been developed for imputing DNAm states for CpG sites in individual cells. For example, Melissa, a Bayesian model, draw inferences solely from DNAm profiles^8^, but it was designed for genomic regions of interest rather than genome-wide imputation. In recent years, deep learning-based models have emerged for imputing single-cell DNAm data, automatically extracting informative features from both DNAm profiles and DNA sequences. One notable model, CpG Transformer^9^, builds upon the transformer model architecture initially designed for language processing. CpG Transformer demonstrated superior performance compared to a previous deep learning model, DeepCpG^10^, which relied on a recurrent neural network. However, both models exhibit limitations in scalability, making them impractical for datasets involving thousands of cells—a scenario increasingly prevalent in the field.

Here, we introduce scMeFormer, a transformer-based and computationally efficient deep learning model, to impute DNAm states for each CpG site in single cells, leveraging information from both local DNA sequences and DNAm profiles across cells. We demonstrate the superior performance of scMeFormer compared to alternative models across four single-nucleus DNAm datasets, each consisting of thousands of nuclei collected from the human brain using distinct technologies^4–7^. Remarkably, scMeFormer exhibits high fidelity in imputing DNAm states, even under significantly reduced coverage—down to 10% of the original CpG sites through downsampling— as demonstrated by its ability to recover cell type clusters and cell type-specific differentially methylated regions (DMRs). We further applied scMeFormer to a single-nucleus DNAm dataset generated from the prefrontal cortex of four schizophrenia patients and neurotypical controls. This enabled the identification of thousands of DMRs associated with schizophrenia that would have remained undetectable without imputation, and added granularity to our understanding of epigenetic alterations in schizophrenia within specific cell types.

## Results

### scMeFormer predicts DNAm states of CpG sites in single cells

The scMeFormer model architecture comprises two main modules: a DNA module and a CpG module (**Figure 1**). The DNA module is designed to learn DNAm motifs, while the CpG module aims to leverage DNAm information from neighboring CpG sites across cell types. The DNA module contains a convolutional neural network (CNN) block for extracting local DNA features, followed by eight layers of transformer blocks to detect distant features that may cooperate to influence DNAm. The CpG module consists of eight layers of transformer blocks that learn features from neighboring CpG sites across cell types. The features learned from both modules are catenated within a fully connected network to predict DNAm states for a given CpG site in a subset of cells whose DNAm states for the given CpG site are measured. The model is trained by minimizing the prediction error, with only measured CpG sites contributing to the loss function in each training step.

**Figure 1.**
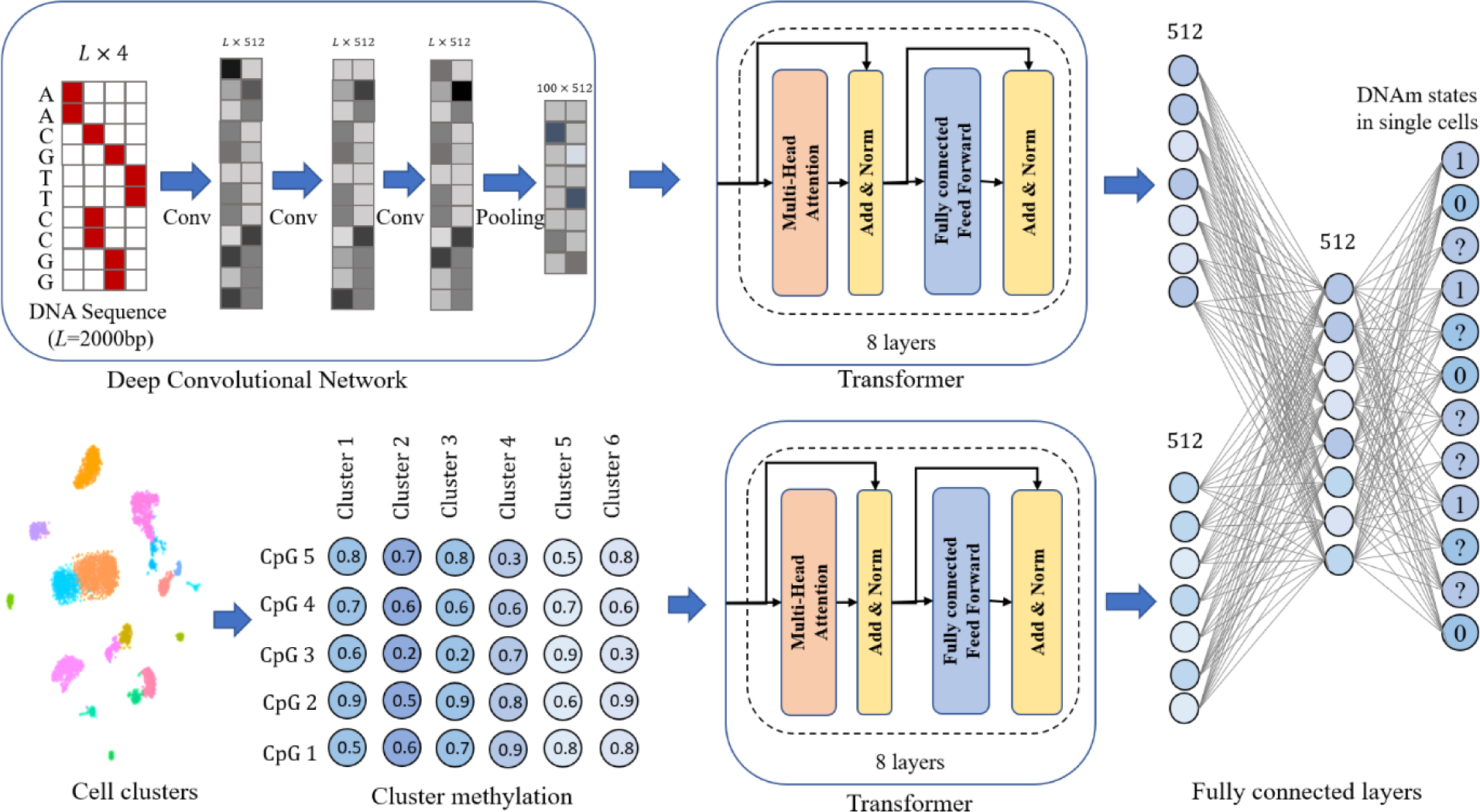
Illustration of scMeformer architecture.

We applied scMeFormer to each of the four single-nucleus DNAm datasets collected from the human brain through four distinct technologies (snmC-seq^4^, snmC-seq2^6^, sn-m3C-seq^5^, and snmCAT-seq^7^). These datasets ranged in size from 2,784 nuclei (snmC-seq) to 4,357 nuclei (snmCAT-seq). Model prediction performance was evaluated on each dataset using independent CpG sites that were not included in both model training and validation. scMeFormer achieved remarkable prediction performance across four datasets, with an average area under the Precision-Recall Curve (AUPRC) of 0.86 (**Figure 2A**). A reduction in prediction performance occurred when scMeFormer employed only the CpG module (average AUPRC = 0.83) or the DNA module (average AUPRC = 0.76), indicating that both modules contribute complementary information for predicting DNAm states. Additionally, we compared scMeFormer with two alternative models: a CNN model and a cluster-based model that imputes CpG states based on the mean DNAm levels of CpG sites in cells of the same cluster. scMeFormer consistently outperformed the two alternative models across the four datasets, except in the snmC-seq dataset, where the cluster model showed slightly better performance (cluster, AUPRC = 0.90; scMeFormer, AUPRC = 0.89) (**Figure 2A**).

**Figure 2.**
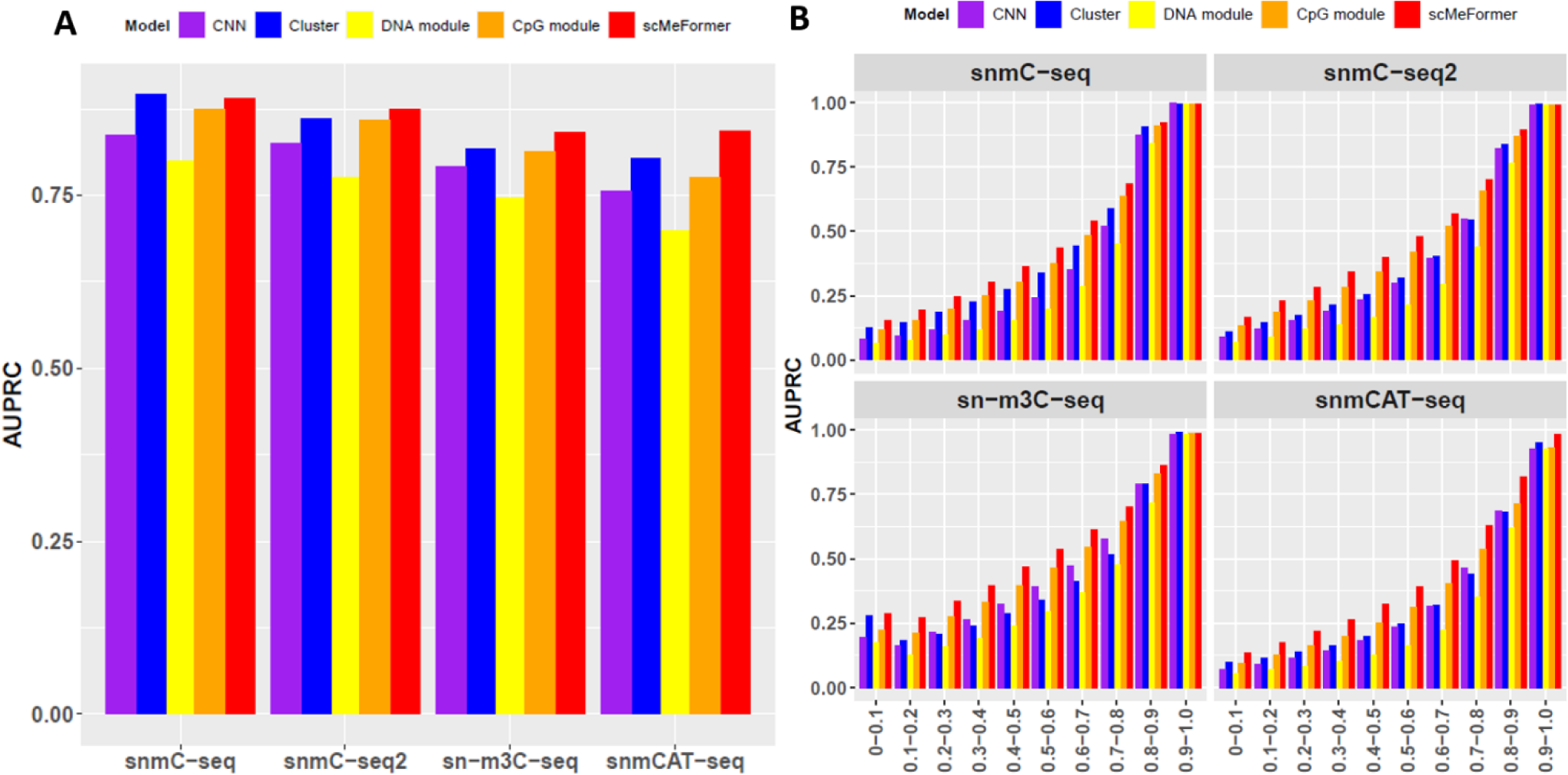
Comparison of model prediction performance between scMeformer and four alternative models across four single-nucleus DNAm datasets. **A.** Comparison was based on all independent testing CpG sites on chromosome 22. **B.** Comparison was based on subsets of independent testing CpG sites stratified by their levels of variations across all cells in each dataset. The “0-0.1” group represents the top 10% variable CpG sites.

We further evaluated the prediction performance of various models on subsets of independent CpG sites, stratified by their levels of variations across cells (**Figure 2B**). All models performed well for CpG sites with the least variability. For example, in three datasets (snmC-seq, snmC-seq2, sn-m3C-seq), all models achieved AUPRC > 0.98 for CpGs ranked in the bottom 10% in terms of their variations. In the snmCAT-seq dataset, scMeFormer achieved a higher AUPRC (0.98) than other models (DNA module: 0.93, CpG module: 0.93, CNN: 0.93, Cluster: 0.95). Notably, we observed a reduction in prediction performance for more variable CpG sites across all models. Nonetheless, scMeFormer consistently outperformed alternative models, particularly for CpG sites in the middle range of variance. For example, within the subset of the most variable CpG sites (ranked in the top 10%), scMeFormer achieved an average AUPRC of 0.19 across four datasets, which was 0.076 and 0.032 higher than the CNN and cluster models, respectively. For CpG sites with intermediate variability (ranked between the top 40% and top 50%), scMeFormer achieved an average AUPRC of 0.40, which was 0.16 higher than the CNN and 0.14 higher than the cluster model across four datasets.

### scMeFormer performance under lower CpG coverage through downsampling

We assessed the performance of scMeFormer under conditions of lower CpG coverage in single cells through downsampling. We systemically reduced the number of CpG sites from 50% to 1% used for training the model to mimic scenarios with lower coverage. In this investigation, we compared scMeFormer with only the cluster model, as the cluster model performed better than CNN in datasets without downsampling. We observed a trend of slightly reduced prediction performance under situations of lower CpG coverage for both models, but scMeFormer consistently outperformed the clustering model across all four datasets, except in the snmC-seq dataset, where the clustering model showed a slightly better performance when the downsampling rate was larger than 0.05 (**Figure 3**). Notably, both models achieved good performance even at 1% of the original CpG coverage (scMeFormer, average AUPRC=0.79; cluster average AUPRC=0.75) (**Figure 3**). We also evaluated the two models under downsampling situations for subsets of independent CpG sites, stratified by their levels of variations across cells (**Supplementary Figure 1**). Similar to our observations for datasets without downsampling, a reduction in prediction performance occurred for more variable CpG sites for both models. However, scMeFormer demonstrated superior performance compared to the cluster model across all four datasets, particularly for CpG sites with intermediate variability. To further assess the fidelity of imputed data under lower CpG coverage, we investigated the ability of imputed CpG sites to recover cell type clusters obtained from the original data. We used the Adjusted Rand Index (ARI) to measure the similarity between clusters obtained from the original data and those derived from: 1) downsampled data without imputation, 2) downsampled data imputed by the cluster model, and 3) downsampled data imputed by scMeFormer. With a predefined cluster number of 10, scMeFormer demonstrated robust performance (ARI > 0.7) across all four datasets even when the downsampling rate was as low as 0.1, but this was not the case for the cluster model and the method without imputation (**Figure 4**). Specifically, scMeFormer achieved an average ARI of 0.88 across the four datasets, whereas the other two methods yielded much lower ARIs (without imputation, ARI=0.50; clustering, ARI=0.51). As downsampling rates became even smaller (< 0.1), all methods exhibited a decline in performance, but scMeFormer showed a slower decay in performance compared to the other two methods (**Figure 4**). This trend remained consistent when increasing the number of predefined clusters (cluster number = 12, 16, 21) (**Supplementary Figure 2**).

**Figure 3.**
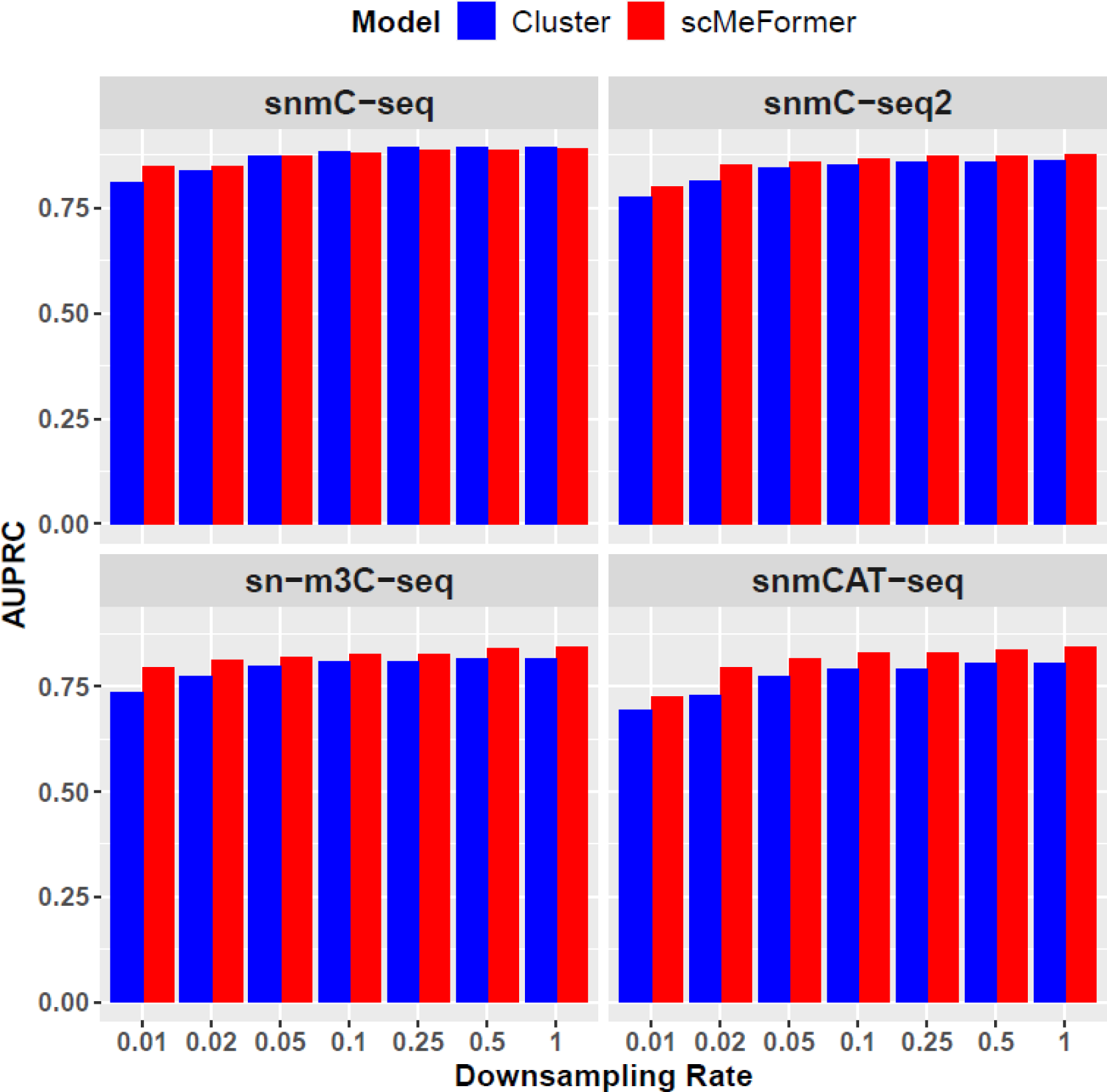
Comparison of model prediction performance between scMeformer and the cluster model across four datasets under lower CpG coverage through downsampling. Comparison was based on independent testing CpG sites on chromosome 22 in each dataset.

**Figure 4.**
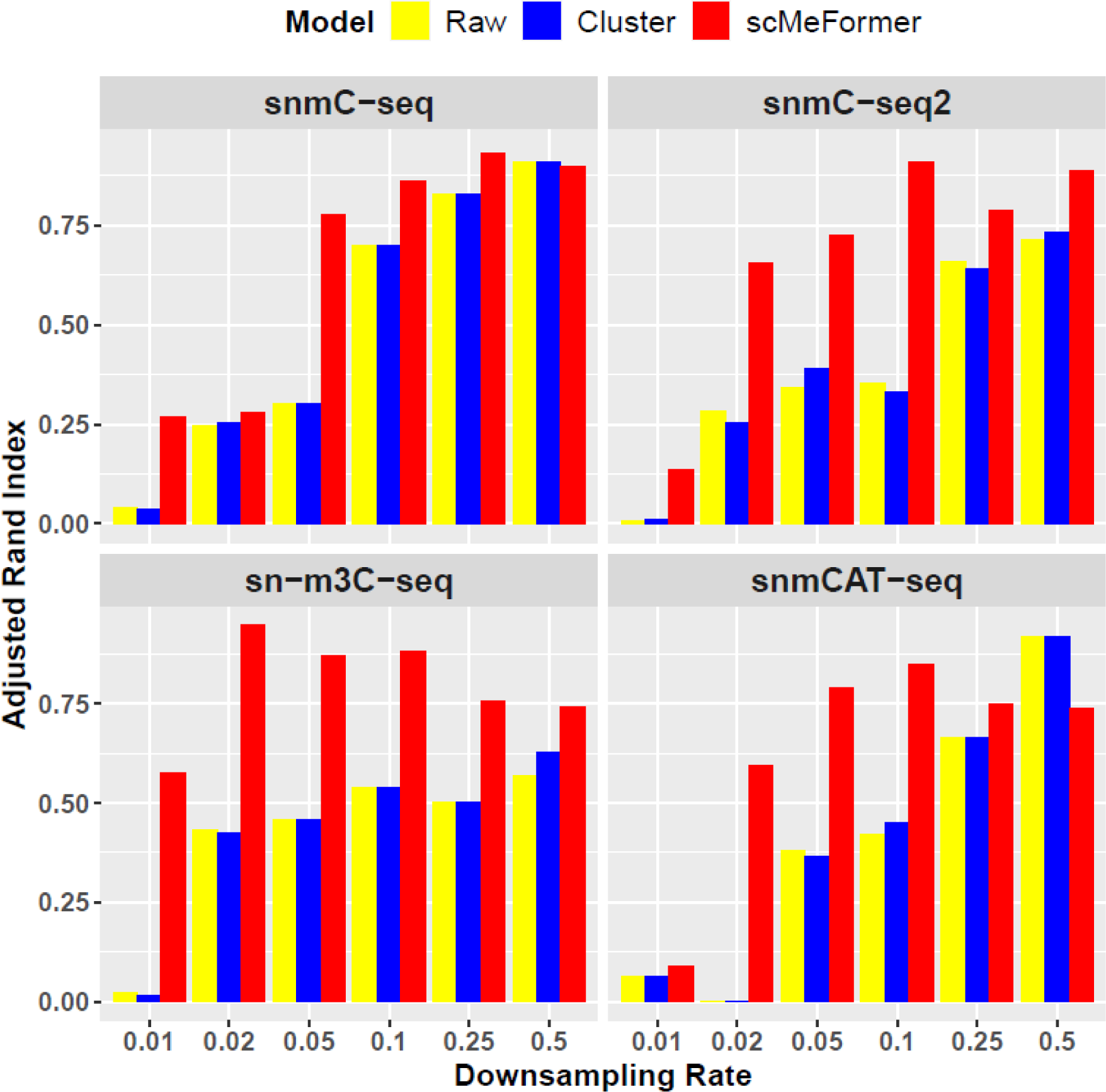
Evaluation of imputed data under lower CpG coverage in its ability to recover cell types identified in the original data across four datasets. The x-axis represents various downsampling rates and the y-axis represents Adjusted Rand Index.

### scMeFormer boosts the detection of cell type-specific DMRs

To further evaluate the quality of imputed data, we assessed its ability to recover DMRs between cell types identified in the original data for two datasets (sn-m3C-seq and snmCAT-seq). We defined cell type-specific DMRs as regions containing at least two differentially methylated CpG sites (DMSs), with each DMS exhibiting lower DNAm levels in the specific cell type, since hypo-DMRs are strong indicators of regulatory elements^11–13^. Remarkably, scMeFormer achieved a high recall rate for cell type-specific DMRs across all cell type pairs in each dataset (sn-m3C-seq: 0.92; snmCAT-seq: 0.92). Even when applied to 10% downsampled data, scMeFormer maintained a high recall rate (sn-m3C-seq: 0.88; snmCAT-seq: 0.86). **Figure 5** illustrates the recall rate for cell type-specific DMRs for each pair of cell types when imputation was performed on the raw and 10% downsampled datasets. This trend remained consistent when employing a stricter definition of cell type-specific DMRs requiring at least five DMSs (**Supplementary Figure 3**). These results demonstrate the efficacy of scMeFormer in imputing CpG sites that faithfully retain information crucial for identifying cell type-specific DMRs.

**Figure 5.**
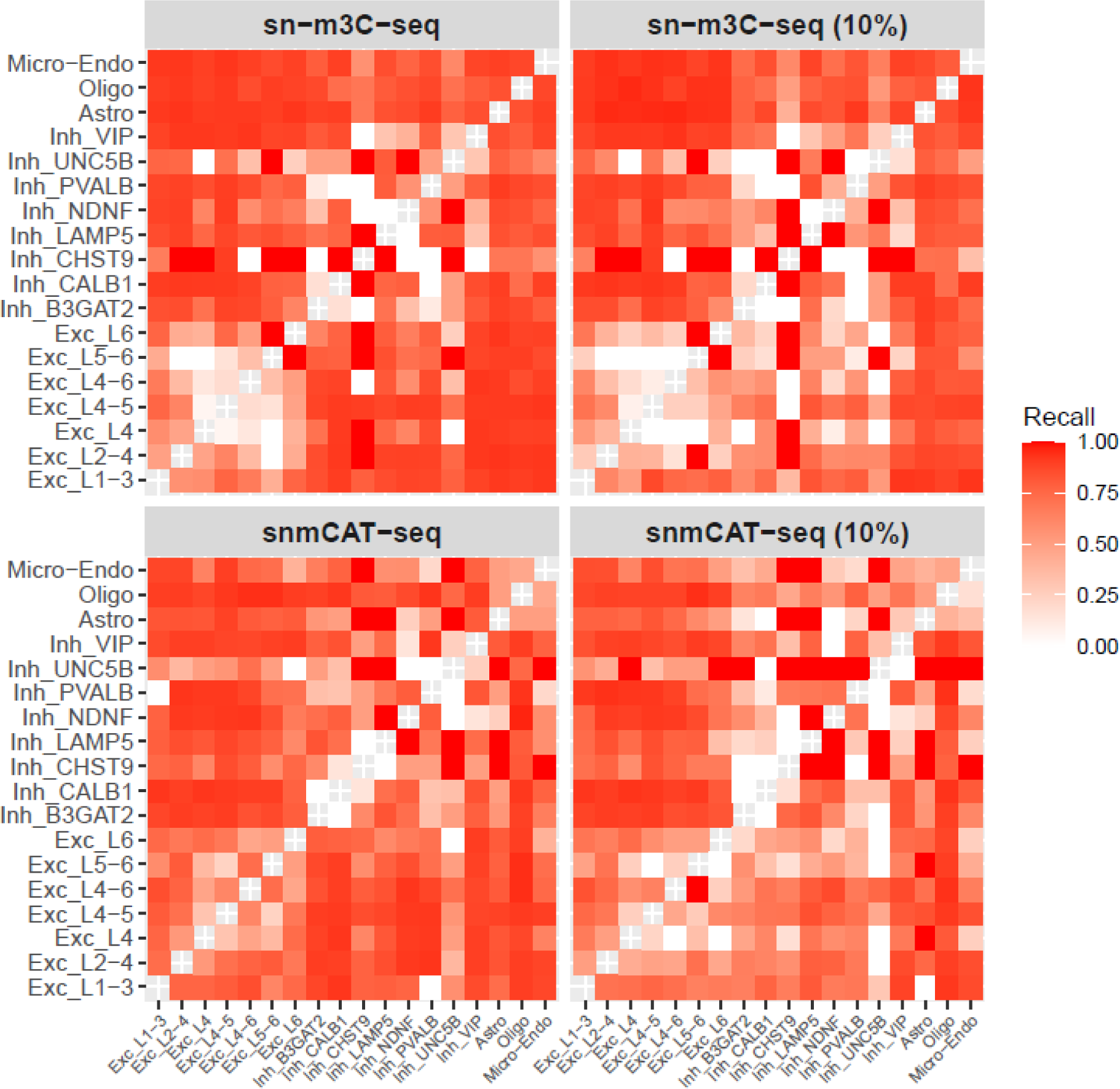
Recall rate for cell type-specific DMRs detected in each pair of cell type in two datasets (sn−m3C−seq and snmCAT−seq). Cell type-specific DMRs were defined by at least two DMSs. The left and right panel shows recall rates when imputation was performed on the raw and 10% downsampled data, respectively. The x-axis represents DMRs specific to each cell type, compared to each cell type labeled on the y-axis.

We compared the numbers of cell type-specific DMRs detected from imputed data with those from raw data. When DMRs were defined by at least two DMSs, we identified an average of 995,342 and 937,227 DMRs across all cell type pairs for the sn-m3C-seq (**Figure 6A**) and snmCAT-seq (**Supplementary Figure 4A**) datasets, respectively. While these numbers were slightly smaller when imputation was applied to the 10% downsampled data (sn-m3C-seq, 933,619 (**Figure 6B**); snmCAT-seq, 944,680 (**Supplementary Figure 4B**)), they remained substantially higher than those identified from raw data (sn-m3C-seq: 634; snmCAT-seq: 145). Additionally, we compared the number of DMRs defined by varying numbers of DMSs, and in general, we observed a smaller number of DMRs with an increasing number of DMSs when scMeFormer was applied to the raw data (**Figure 6A, Supplementary Figure 4A**). These numbers became slightly smaller but remained comparable when scMeFormer was applied to the 10% downsampled data (**Figure 6B, Supplementary Figure 4B**).

**Figure 6.**
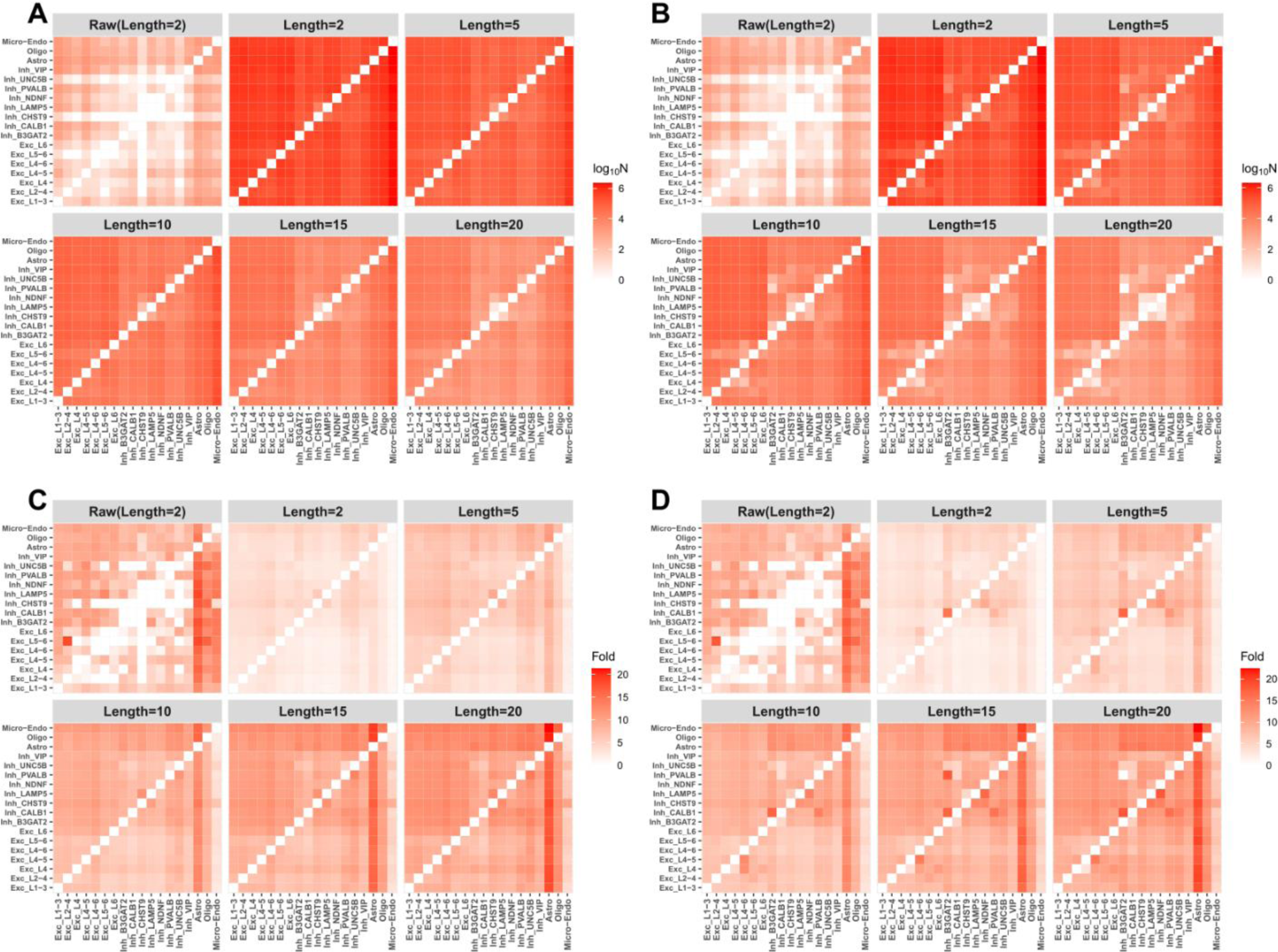
scMeFormer enhances the detection of cell type-specific DMRs in the dataset of sn-m3C-seq. **A**. The top left panel displays the number of cell type-specific DMRs (log scale) detected in the raw data, with subsequent panels showing the number of cell type-specific DMRs (log scale) detected in imputed data, defined by varying numbers of DMSs. The x-axis represents DMRs specific to each cell type, compared to each cell type labeled on the y-axis. **B**. Similar to A, but imputation was performed on 10% downsampled data. **C**. Enrichment fold for H3K27ac mark in the corresponding broad cell type among cell type-specific DMRs detected in the raw data (top left panel) and in the imputed data defined by varying number of DMS. The x-axis represents DMRs specific to each cell type, compared to each cell type labeled on the y-axis. **D**. Similar to C, but imputation was performed on 10% downsampled data.

To estimate the false positive rates of our model in calling cell type-specific DMRs, we turned to two broad cell types (neuron and glia) in the sn-m3C-seq dataset that allows us to identify CpG sites of sufficient coverage for identifying differentially methylation CpG sites in the raw data. Specifically, we first collected 7,512 CpG sites not within blacklist regions^14^ but with high coverage (≥ 1000) in each broad cell type. Among these, 1,739 showed differential methylation between the two cell types (FDR < 0.05). We then ran DMR analysis between the two cell types in imputed data, and selected a subset of DMRs (1,288) that contained as least one CpG site with coverage ≥ 1000 in each broad cell type in the raw data. Among the 1,131 DMRs we examined, 534 contained at least one differentially methylated CpG site from the selected high-coverage CpG sites in the raw data, suggesting that 47% of them are likely true DMRs (precision = 0.47). When we extended the DMRs by a window of 1kb, we observed 958 DMRs contained at least one differentially methylated CpG site of high coverage in the raw data, resulting in a precision of 0.85.

To assess the biological relevance of cell type-specific DMRs identified from imputed data, we leveraged histone mark (H3K27ac) data that indicates active enhancers in four broad cell types (neuron, astrocyte, oligo, and microglia)^15^. Given prior evidence that regulatory regions are associated with low DNAm levels, we hypothesized that reliable cell type-specific DMRs should be enriched for H3K27ac marks in the corresponding broad cell type. Our analysis confirmed this hypothesis, revealing that cell type-specific DMRs were enriched for H3K27ac-marked regions in the corresponding broad cell type across both datasets (**Figure 6C, Supplementary Figure 4C**). Remarkably, this enrichment pattern remained consistent even when scMeFormer was applied to the 10% downsampled data (**Figure 6D, Supplementary Figure 4D**). This enrichment pattern became stronger when DMRs were defined by a larger number of DMSs, suggesting a higher probability of these regions being regulatory.

To further validate the quality of cell type-specific DMRs identified from imputed data, we examined their enrichment for the heritability of three brain disorders (schizophrenia, bipolar, and depression) and human height. This evaluation was based on the rationale that if cell type-specific DMRs were enriched for regulatory regions specific to those cell types, they would also enrich the heritability of traits in cell types relevant to the traits. This was indeed the case for DMRs defined by at least five CpGs in the sn-m3C-seq dataset (**Figure 7**). We observed strong enrichment for all three brain disorders, but not height, for DMRs specific to excitatory neuronal cell types, particularly when they were compared to inhibitory or non-neuronal cell types. Enrichment was also observed for DMRs specific to inhibitory neuronal cell types when they were compared to non-neuronal cell types. Conversely, no enrichment was observed for DMRs specific to non-neuronal cell types across all three brain disorders, except for depression, where we noted enrichment for astrocyte-specific DMRs, supporting the emerging role of this cell type in depression^16^. This pattern remained consistent for DMRs defined by at least 10 DMSs and DMRs (defined by at least 5 or 10 DMSs) detected from 10% downsampled data (**Supplementary Figure 5**). Heritability enrichment analysis for DMRs detected in the snmCAT-seq dataset showed similar pattern (**Supplementary Figure 6**).

**Figure 7.**
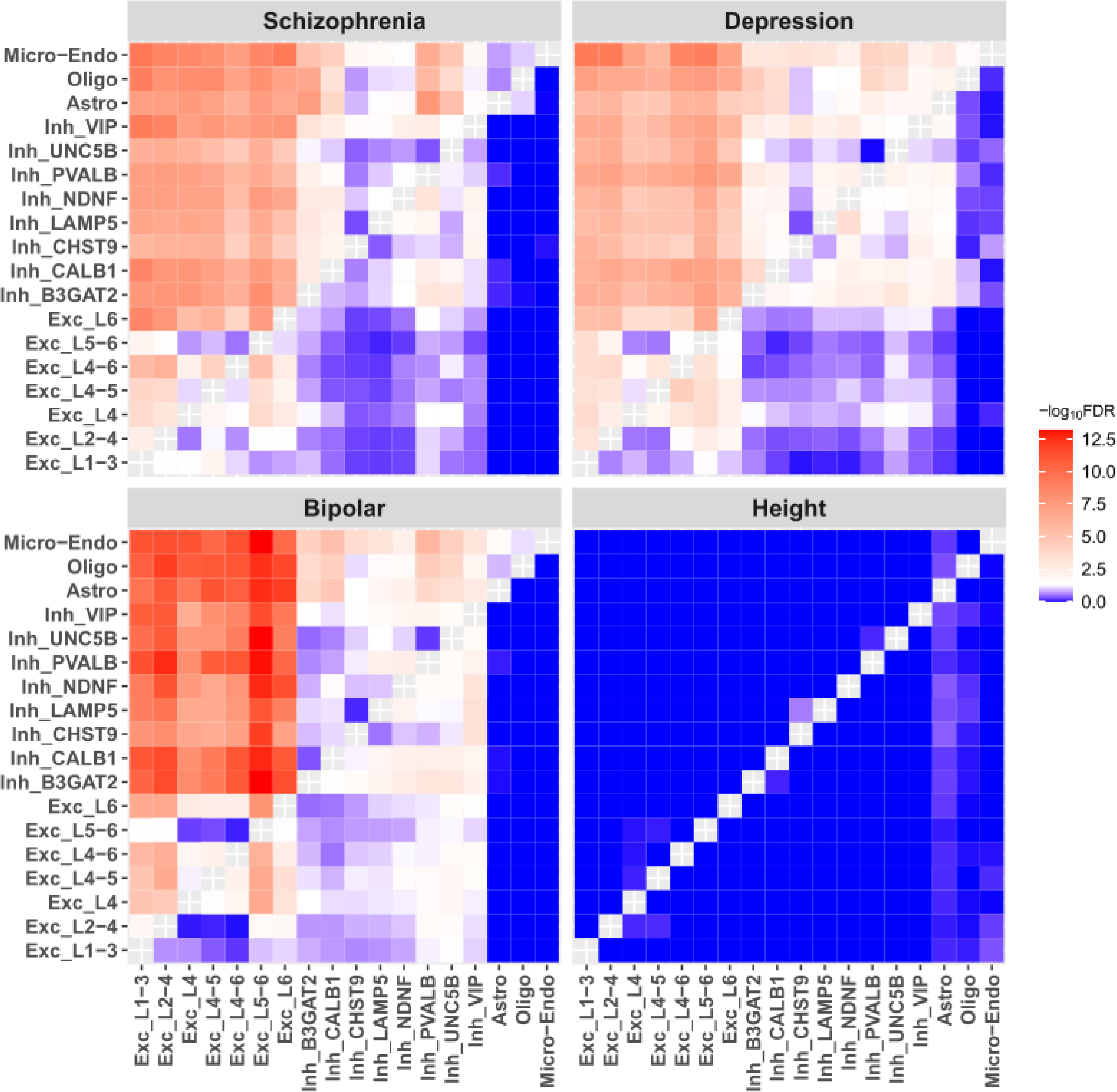
Heritability enrichment analysis for cell type-specific DMRs identified from imputed data of the sn-m3C-seq dataset. The x-axis represents DMRs specific to each cell type, compared to each cell type labeled on the y-axis. Cell type-specific DMRs were defined by at least five DMSs. The color represents the −log_10_(FDR) values, derived from the z-score of per-SNP heritability as reported by stratified LDSC regression.

### scMeFormer enhances the detection of schizophrenia-associated DMRs

We applied scMeFormer to impute CpG states in 2,534 single-nucleus DNAm profiles generated from the prefrontal cortex of four schizophrenia cases and four neurotypical controls. We first evaluated whether imputed data could recover clusters derived from CpHs, which is better suited for clustering neuronal cell types than CpGs^4^. Specifically, using DNAm levels of CpHs in nonoverlapping 100kb bins, we first clustered these nuclei into five major cell types: two excitatory neuron subtypes from the superficial (SupExc) and deep cortical layers (DeepExc), two inhibitory neuron subtypes from the cortical ganglionic eminence (InhCGE) and the medial ganglionic eminence (InhMGE), and a glial cell type. We then clustered the same set of nuclei using imputed DNAm levels of CpGs in nonoverlapping 100kb bins. Clusters from imputed data closely mirror the clusters obtained using CpHs (**Supplementary Figure 7**), indicating that the imputation process preserved data properties related to cell type composition.

Next, we aimed to identify schizophrenia-associated DMRs within each cell type. Notably, no DMRs were detected before imputation, but imputation revealed a substantial number of DMRs across the five cell types (**Figure 8A**). S-LDSC analysis demonstrated that cell type-specific DMRs were enriched for schizophrenia heritability, particularly in neuronal cell types and, to a lesser extent, in glial cells (**Figure 8B**). While enrichment was also observed for bipolar disorder and depression, the strength was generally weaker compared to schizophrenia. There was no enrichment observed for height heritability, suggesting these DMRs are relatively specific to schizophrenia. Furthermore, we linked the DMRs detected in each cell type to the genes they may regulate, leveraging chromatin loops detected in the broad neuron and glial cell types in a previous study^15^. We observed that DMRs in deep excitatory neurons were linked to genes enriched for neurodevelopmental processes, whereas DMRs in super excitatory neurons were linked to genes enriched for synaptic signaling (**Figure 8C**). While genes regulated by DMRs in inhibitory neurons did not reveal significant pathways, the top pathways were consistent with those from DMRs in excitatory neurons. Interestingly, the top enriched pathways for genes regulated by DMRs in the glial cell type were related to the immune system, although not statistically significant after FDR correction (FDR > 0.05). These findings add granularity to our understanding of the functional implications of these epigenetic alterations in schizophrenia within specific cell types.

**Figure 8.**
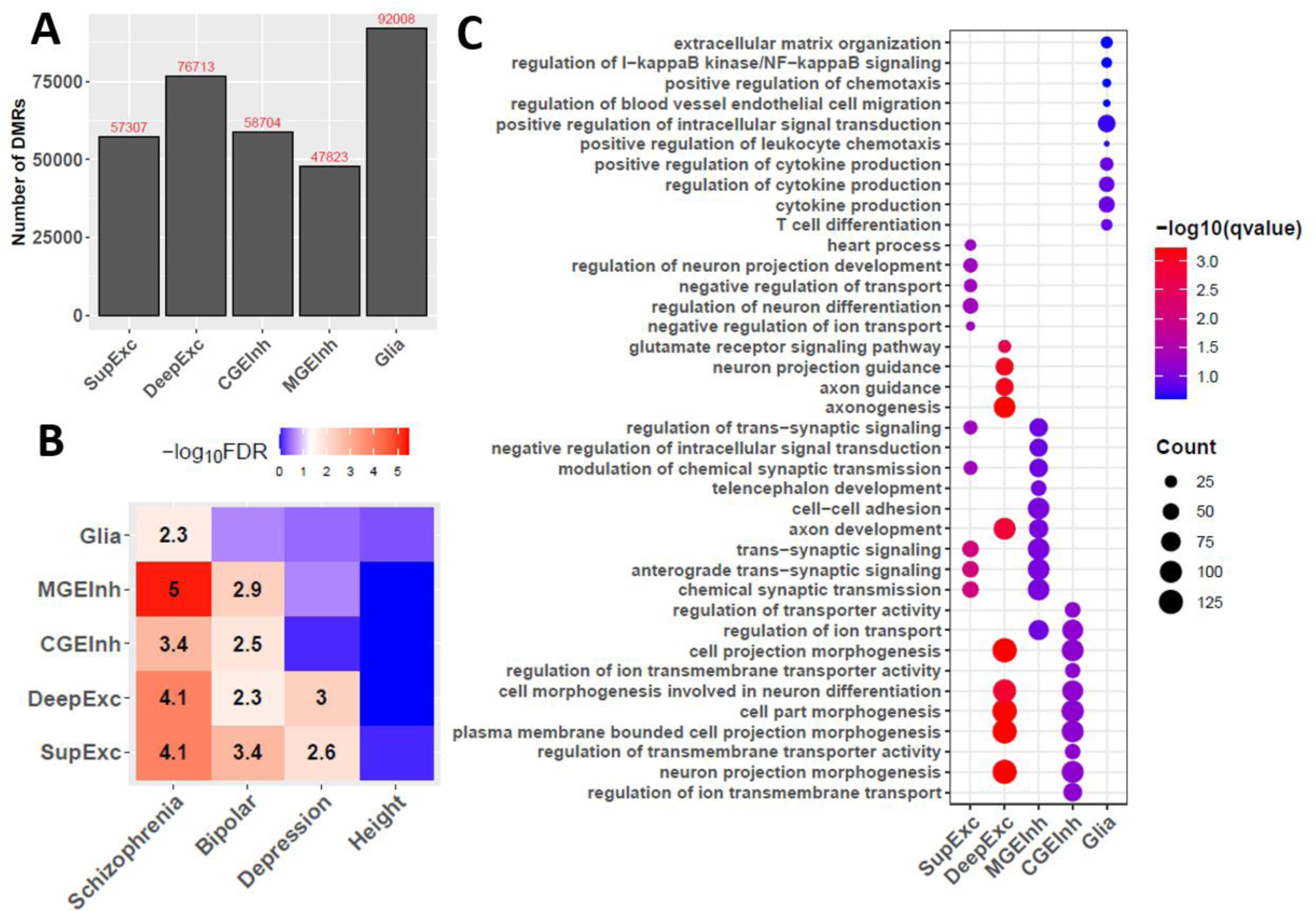
scMeFormer enhances the detection of schizophrenia-associated DMRs. **A**. Numbers of schizophrenia-associated DMRs detected in each cell type. **B**. Evaluation of schizophrenia-associated DMRs for their contributions to the heritability of three brain disorders and human height. The numbers within the squares are z-scores of per-SNP heritability that are significant after multiple testing correction (FDR < 0.05). **C**. Top 10 enriched pathways for genes linked by schizophrenia-associated DMRs through chromatin loops in each cell type.

## Discussion

Single-cell DNAm profiling technologies have offered unprecedented opportunities to explore the epigenetic landscape of DNA sequence at the single-cell resolution. However, many of these technologies suffer from a high missing rate of CpG sites, limiting their full potential to uncover the epigenetic mechanism underlying various biological processes and diseases. Previous deep learning models have attempted to impute CpGs methylation status in single cells^9, 10^. However, these models face challenges in scaling to thousands of cells, a scenario that is becoming increasingly popular. In contrast, our novel model, scMeFormer, efficiently imputes data for thousands of cells. It achieves training times of approximately 72 hours for each single-nucleus DNAm dataset we investigated, utilizing four NVIDIA A100 GPUs. Crucially, the multitask prediction framework employed by scMeFormer allows the imputation of single-cell DNAm datasets of any size without incurring additional computational time costs. Remarkably, scMeFormer exhibits the ability to impute DNAm states with high fidelity, even with only one-tenth of the current coverage of CpG sites through downsampling, as evidenced by the recovery of cell type clusters and cell type-specific DMRs. Furthermore, we applied scMeFormer to a single-nucleus DNAm dataset generated from the prefrontal cortex of schizophrenia patients and neurotypical controls. This led to the identification of thousands of schizophrenia-associated DMRs that would have remained undetectable without imputation, adding granularity to our understanding of epigenetic alterations in schizophrenia within specific brain cell types. Our study underscores the power of deep learning in imputing DNA states in single cells, and we expect that scMeFormer will be a valuable tool for single-cell DNAm studies.

While scMeFormer demonstrates significant promise for single-cell DNAm data imputation, we acknowledge several limitations and potential areas for future work. First, scMeFormer relies on input DNA sequences from the reference genome, which does not precisely align with the DNA sequences in the samples we used for training the model. The model’s performance could be further improved by utilizing DNA sequences and DNAm data from the same individuals. Second, scMeFormer currently focuses on CpG sites imputation. Expanding its capabilities to CpH sites, known for their crucial roles in neurons, would enhance its utility. Third, scMeFormer requires retraining for each new dataset. Building a transferable model capable of adapting to new datasets without retraining would be highly beneficial for broader applications.

## Methods

### Training datasets

Training datasets include four single-nucleus DNAm datasets generated by four different technologies we developed: snmC-seq^4^, snmC-seq2^6^, sn-m3C-seq^5^, and snmCAT-seq^7^, all applied to human postmortem brain tissues. Detailed information regarding each technology and the bioinformatics procedures for data processing have been described in our original studies. Briefly, snmC-seq was a multiplexed single-nucleus DNAm profiling technique we initially developed, which was used to analyze the methylomes of 2,784 neurons isolated from the human frontal cortex. snmC-seq2 was the improved version of snmC-seq with increased read mapping and enhanced throughput, and was used to profile the methylomes of 3,072 nuclei obtained from postmortem prefrontal cortex. sn-m3C-seq was a single-cell multi-omics technique that jointly profiles chromatin conformation and DNAm from the same cells and was applied to profile 4,237 nuclei from human BA10 cortical tissue. snmCAT-seq was developed to jointly profile methylome, chromatin accessibility, and transcriptome from the same cells, and was applied to profile 4,357 nuclei isolated from postmortem human BA10 cortical tissue.

### scMeformer architecture

scMeformer contains three main modules: a DNA module, a CpG module, and a fully connected network. The input to the model includes two modalities: a one-hot encoded DNA sequence of 2 kb and the DNAm levels of 100 neighboring CpGs around the target CpG in each cell cluster. The output of the model is the predicted methylation states of the target CpG in each cell across all cells. Below are descriptions of each module.

#### DNA module

The DNA module employs the INTERACT architecture developed in our previous study^17^, which consists of two sub-modules: a convolutional neural network (CNN) and the encoder of transformer. The CNN comprises three convolutional layers, a batch normalization layer, a max-pooling layer, and a dropout layer. Each convolutional layer uses 512 kernels of size 10 to learn motifs from DNA sequences, and each kernel is activated by a rectified linear unit (ReLU) function. The max-pooling layer (step size = 20bp) is used after the three convolutional layers to capture motifs learned by the convolutional layers. The batch normalization layer is used after the max-pooling layer to improve training speed and stability. The dropout layer is employed to prevent overfitting. The dropout rate in this layer is set to 0.5.

The encoder of transformer consists of a stack of eight identical layers, which takes the CNN output as input to learn distant features that may act jointly. Each layer in the transformer encoder employs two sub-layers. The first sub-layer is a multi-head self-attention layer that learns the attention between any two features at different positions. The second sub-layer is a simple, position-wise fully connected feed-forward network. After both sub-layers, a normalization layer is employed to speed up training and improve training stability. Additionally, a dropout layer with a rate of 0.1 is employed to prevent overfitting. Each sub-layer in the encoder has a residual connection to help mitigate the vanishing gradient problem. Residual connections are often used in deep neural networks to prevent the network from forgetting important features of the input during training.

#### CpG module

The CpG module employs the same encoder of transformer as the DNA module. Given a target CpG site the model aims to predict, the CpG module takes as input the DNAm levels of 100 CpG sites around the target CpG site (excluding the target CpG site itself) across pre-defined cell clusters. To create this input, all single cells are initially clustered into clusters based on the DNAm levels of CpG sites within non-overlapping 100kb bins. For a given CpG site, its DNAm level in each cluster is determined by dividing the number of methylated reads by the total number of reads in that cluster. If the cells are clustered into *n* clusters, the input to the CpG module is a 100 × n matrix.

#### Fully connected network

The fully connected network is composed of a hidden layer with 512 units, a dropout layer, and an output layer. The dropout layer is designed to prevent overfitting and uses a drop rate of 0.1. The output layer applies the sigmoid function to scale the predicted values between 0 and 1. The number of units in the output layer is equal to the number of cells. The input to the fully connected network is the concatenation of the output from the DNA module and the CpG module. Each unit in the output layer represents the DNAm state of the target CpG site in the corresponding cell and is indicated by a value of either 0 or 1.

#### Model training

We divided CpG sites into three subsets by chromosomes for model training, validation, and evaluation. The training set consisted of CpG sites on chromosomes 1 to 20, while CpG sites on chromosome 21 were used as the validation set for model tuning, and CpG sites on chromosome 22 were used as the independent testing set to evaluate the model prediction performance. Approximately 5% of CpG sites were covered by at least one read in a single cell across the four datasets and were employed for model training, while the remaining CpG sites were not covered by any reads and were not used in model training. We defined DNAm state as 1 for a CpG site if all mapped reads support methylation, and 0 if all mapped reads support unmethylation. We did not include CpG sites for model training if their mapped reads support both methylation and unmethylation.

#### Alternative models

We compared scMeformer to four alternative models: the CNN model, the cluster model, the scMeformer model but with only the DNA module, and the scMeformer model but with only the CpG module. The CNN model also consists of three modules, including a DNA module, a CpG module, and a fully connected network. However, in the CNN model, both the DNA and CpG modules employ a convolutional neural network (the same as described in the DNA module of scMeformer) rather than the transformer encoders. The cluster model first clusters cells into clusters based on DNAm levels of CpGs within non-overlapping 100kb bins. For each cell in a cluster, the methylation state for a CpG site not covered by any reads is imputed by known methylation states of this CpG site in cells of the same cluster.

### Cell type-specific DMRs

We leveraged pre-assigned cell type labels for each nucleus in each dataset from the original studies. We then employed the DMRfind function from methylpy (v1.4.2)^18^ to identify cell type-specific DMRs across all cell type pairs. Briefly, DMRfind utilizes a permutation-based root mean square test of goodness-of-fit to identify differentially methylated sites (DMS) across samples. Consecutive DMSs within 250 bp are then merged into DMRs. In our imputed dataset, we considered a CpG site with a predicted methylation status as covered by one read supporting methylation, and a predicted unmethylated CpG site as covered by one read supporting unmethylation.

### Stratified LD score regression

We performed stratified LD score regression (S-LDSC)^19^ to evaluate the enrichment of heritability of three brain disorders (schizophrenia^20^, depression^21^, and bipolar disorder^22^). We also included one non-brain trait, human height^23^, as a negative control to examine whether our findings are specific to brain disorders. Following recommendations from the LDSC resource website (https://alkesgroup.broadinstitute.org/LDSCORE), S-LDSC was run for each list of variants with the baseline LD model v2.2 that included 97 annotations to control for the LD between variants with other functional annotations in the genome. We used HapMap Project Phase 3 SNPs as regression SNPs, and 1000 Genomes SNPs of European ancestry samples as reference SNPs, which were all downloaded from the LDSC resource website. To evaluate the unique contribution of annotations to trait heritability, we utilized a metric from S-LDSC: the z-score of per-SNP heritability. This metric allows us to discern the unique contributions of candidate annotations while accounting for contributions from other functional annotations in the baseline model. The p-values were derived from the z-score assuming a normal distribution and FDR was computed from the p-values based on Benjamini & Hochberg procedure.

### Single-nucleus DNAm data from schizophrenia cases and controls

We generated single-nucleus methylomes from the prefrontal cortex of four schizophrenia cases and four neurotypical controls using the snmCAT-seq technique, but without capturing chromatin accessibility information. The brain tissues were from the brain repository at the Lieber Institute for Brain Development. Details on tissue acquisition, processing, curation, and dissection, were described in prior reports^24^. All eight brain samples were male Caucasian individuals with a mean age of 42 years old in both cases and controls.

We processed sequencing reads by implementing a versatile mapping pipeline (http://cemba-data.readthedocs.io/) for all the methylome-based technologies developed by our group, as detailed in our previous study^7^. After allc files were generated, the methylcytosine (*mc*) and total cytosine basecalls (*cov*) were summed up for each 100kb bin across the hg19 genome for each sequence context (CG, CH). After filtering cells by various mapping metrics, 2,534 nuclei were retained for further analysis. Using DNAm levels of CpHs in nonoverlapping 100kb bins, we clustered these nuclei into five major cell types: two excitatory neuron subtypes from the superficial and deep cortical layers (SupExc and DeepExc), two inhibitory neuron subtypes from the cortical ganglionic eminence (InhCGE) and the medial ganglionic eminence (InhMGE), and a glial cell type. We also clustered these nuclei using DNAm levels of CpGs in nonoverlapping 100kb bins after imputation employing functions from the scanorama package. We employed the DMRfind function from methylpy to identify schizophrenia-associated DMRs for each cell type. DMRs were called for regions with at least two DMSs (FDR < 0.01) within 250bp and each DMS had the same direction of effect in at least two samples from either the case or control group. DMR detected in neuronal and glial cell types were assigned to target genes, leveraging reported chromatin loops between active promoters and distal regulatory regions in the broad neuronal and two glial cell types (microglia and oligodendrocytes)^15^. We used clusterProfiler^25^ for gene ontology enrichment analysis for genes regulated by DMRs detected in each cell type, using all distally regulated genes detected in the corresponding broad cell type as the background genes.

## Acknowledgments

We are grateful for the vision and generosity of the Lieber and Maltz families, who made this work possible; the families who donated to this research; Computing support from The Joint High Performance Computing Exchange (JHPCE) facility in the Department of Biostatistics at the Johns Hopkins Bloomberg School of Public Health. This study was supported by National Institutes of Health grants R01MH121394 and R01MH112751 (to S.H.).

## Conflict of interest

DRW serves on the Scientific Advisory Boards of Sage Therapeutics and Pasithea Therapeutics. JRE serves on the scientific advisory board of Zymo Research.

---

**Supplementary Figure 1.** Comparison of prediction performance between scMeformer and the cluster model across four datasets under lower CpG coverage through downsampling. Comparison was based on subsets of independent testing CpG sites stratified by their levels of variation across cells. The “0-0.1” group represents the top 10% variable CpG sites.

**Supplementary Figure 2.** Evaluation of imputed data quality under reduced CpG coverage in recovering cell types identified in the original data across four datasets, when varying the number of defined clusters in the original data from 12 to 21.

**Supplementary Figure 3.** Recall rate for cell type-specific DMRs for all cell type pairs in two datasets (sn−m3C−seq and sn−m3C−seq). Cell type-specific DMRs were defined by at least five DMSs. The left and right panel shows recall rate when imputation was conducted on the raw and 10% downsampled data, respectively. The x-axis represents DMRs specific to each cell type, compared to each cell type labeled on the y-axis.

**Supplementary Figure 4.** scMeFormer enhances the detection of cell type-specific DMRs in the dataset of snmCAT-seq. **A**. The top left panel displays the number of cell type-specific DMRs (log scale) detected in the raw data, with subsequent panels showing the number of cell type-specific DMRs (log scale) detected in imputed data, defined by varying numbers of DMSs. The x-axis represents DMRs specific to each cell type, compared to each cell type labeled on the y-axis. **B**. Similar to A, but imputation was performed on 10% downsampled data. **C**. Enrichment fold for H3K27ac mark in the corresponding broad cell type among cell type-specific DMRs detected in the raw data (top left panel) and in the imputed data defined by varying number of DMS. The x-axis represents DMRs specific to each cell type, compared to each cell type labeled on the y-axis. **D**. Similar to C, but imputation was performed on 10% downsampled data.

**Supplementary Figure 5.** Heritability enrichment analysis for cell type-specific DMRs identified from imputed data in the dataset of sn-m3C-seq. These plots show enrichment analysis results for cell type-specific DMRs defined by at least five or 10 DMSs and when imputation was performed in original data or 10% downsampled data. The color represents the −log_10_(FDR) values, derived from the z-score of per-SNP heritability as reported by stratified LDSC regression. The x-axis represents DMRs specific to each cell type, compared to each cell type labeled on the y-axis.

**Supplementary Figure 6.** Heritability enrichment analysis for cell type-specific DMRs identified from imputed data in the dataset of snmCAT-seq. These plots show enrichment analysis results for cell type-specific DMRs defined by at least five or 10 DMSs and when imputation was performed in original data or 10% downsampled data. The color represents the −log_10_(FDR) values, derived from the z-score of per-SNP heritability as reported by stratified LDSC regression. The x-axis represents DMRs specific to each cell type, compared to each cell type labeled on the y-axis.

**Supplementary Figure 7.** T-SNE plot using imputed CpG sites resemble clusters obtained using CpHs in the raw data.

## References

1. Greenberg, M.V.C. & Bourc’his, D. The diverse roles of DNA methylation in mammalian development and disease. Nat Rev Mol Cell Biol 20, 590–607 (2019).

2. Ahn, J., Heo, S., Lee, J. & Bang, D. Introduction to Single-Cell DNA Methylation Profiling Methods. Biomolecules 11 (2021).

3. Luo, C., Hajkova, P. & Ecker, J.R. Dynamic DNA methylation: In the right place at the right time. Science 361, 1336–1340 (2018).

4. Luo, C. et al. Single-cell methylomes identify neuronal subtypes and regulatory elements in mammalian cortex. Science 357, 600–604 (2017).

5. Lee, D.S. et al. Simultaneous profiling of 3D genome structure and DNA methylation in single human cells. Nat Methods 16, 999–1006 (2019).

6. Luo, C. et al. Robust single-cell DNA methylome profiling with snmC-seq2. Nat Commun 9, 3824 (2018).

7. Luo, C. et al. Single nucleus multi-omics identifies human cortical cell regulatory genome diversity. Cell Genom 2 (2022).

8. Kapourani, C.A. & Sanguinetti, G. Melissa: Bayesian clustering and imputation of single-cell methylomes. Genome Biol 20, 61 (2019).

9. De Waele, G., Clauwaert, J., Menschaert, G. & Waegeman, W. CpG Transformer for imputation of single-cell methylomes. Bioinformatics 38, 597–603 (2022).

10. Angermueller, C., Lee, H.J., Reik, W. & Stegle, O. DeepCpG: accurate prediction of single-cell DNA methylation states using deep learning. Genome Biol 18, 67 (2017).

11. Stadler, M.B. et al. DNA-binding factors shape the mouse methylome at distal regulatory regions. Nature 480, 490–495 (2011).

12. Hon, G.C. et al. Epigenetic memory at embryonic enhancers identified in DNA methylation maps from adult mouse tissues. Nat Genet 45, 1198–1206 (2013).

13. Mo, A. et al. Epigenomic Signatures of Neuronal Diversity in the Mammalian Brain. Neuron 86, 1369–1384 (2015).

14. Amemiya, H.M., Kundaje, A. & Boyle, A.P. The ENCODE Blacklist: Identification of Problematic Regions of the Genome. Sci Rep 9, 9354 (2019).

15. Nott, A. et al. Brain cell type-specific enhancer-promoter interactome maps and disease-risk association. Science 366, 1134–1139 (2019).

16. O’Leary, L.A. & Mechawar, N. Implication of cerebral astrocytes in major depression: A review of fine neuroanatomical evidence in humans. Glia 69, 2077–2099 (2021).

17. Zhou, J. et al. Deep learning predicts DNA methylation regulatory variants in the human brain and elucidates the genetics of psychiatric disorders. Proc Natl Acad Sci U S A 119, e2206069119 (2022).

18. Schultz, M.D. et al. Human body epigenome maps reveal noncanonical DNA methylation variation. Nature 523, 212–216 (2015).

19. Finucane, H.K. et al. Partitioning heritability by functional annotation using genome-wide association summary statistics. Nat Genet 47, 1228–1235 (2015).

20. Trubetskoy, V. et al. Mapping genomic loci implicates genes and synaptic biology in schizophrenia. Nature 604, 502–508 (2022).

21. Howard, D.M. et al. Genome-wide meta-analysis of depression identifies 102 independent variants and highlights the importance of the prefrontal brain regions. Nat Neurosci 22, 343–352 (2019).

22. Mullins, N. et al. Genome-wide association study of more than 40,000 bipolar disorder cases provides new insights into the underlying biology. Nat Genet 53, 817–829 (2021).

23. Yengo, L. et al. Meta-analysis of genome-wide association studies for height and body mass index in approximately 700000 individuals of European ancestry. Hum Mol Genet 27, 3641–3649 (2018).

24. Jaffe, A.E. et al. Mapping DNA methylation across development, genotype and schizophrenia in the human frontal cortex. Nat Neurosci 19, 40–47 (2016).

25. Yu, G., Wang, L.G., Han, Y. & He, Q.Y. clusterProfiler: an R package for comparing biological themes among gene clusters. OMICS 16, 284–287 (2012).

