## Supplementary figures and images for "scMeFormer: a transformer-based deep learning model for imputing DNA methylation states in single cells enhances the detection of epigenetic alterations in schizophrenia"

### Supplementary Figure 1

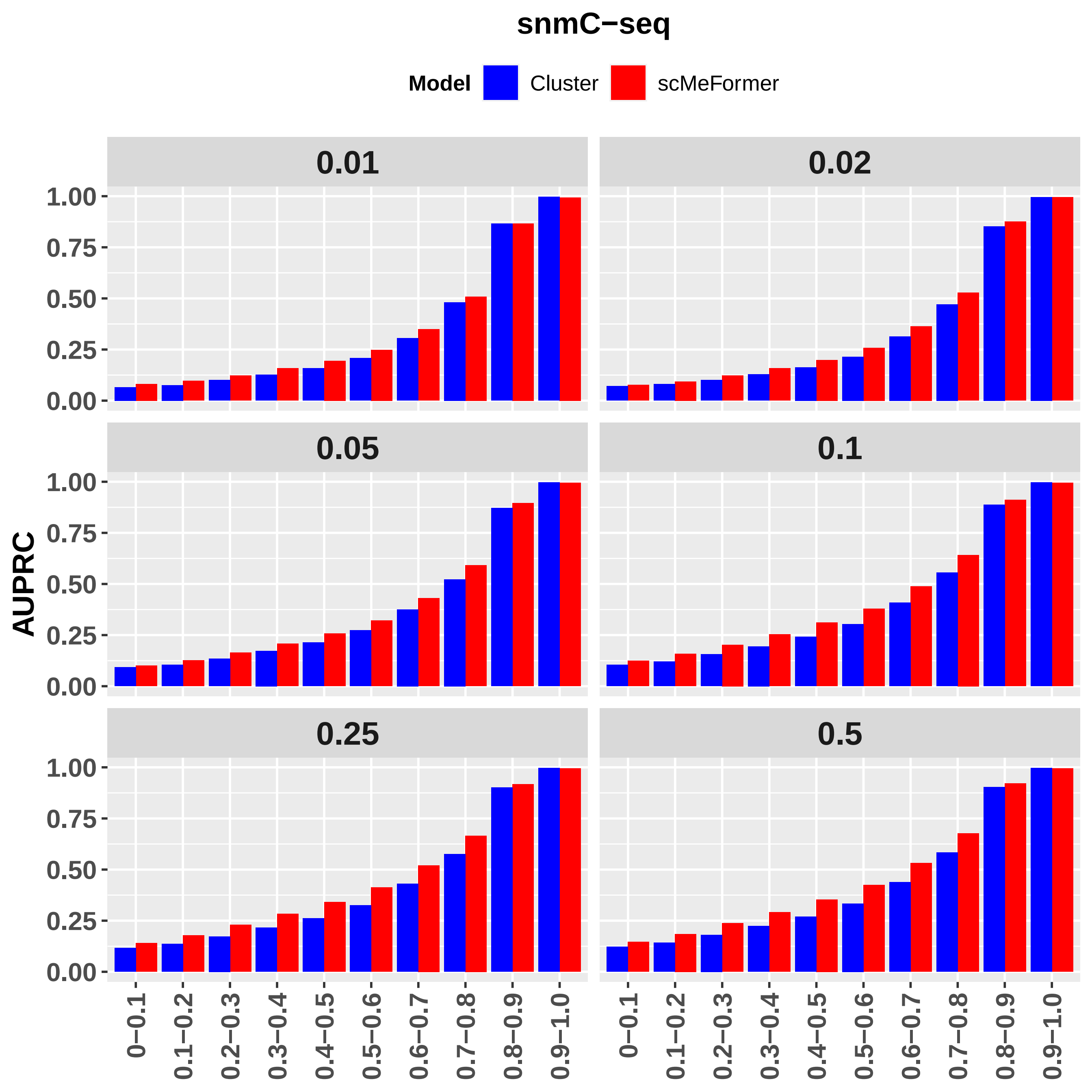

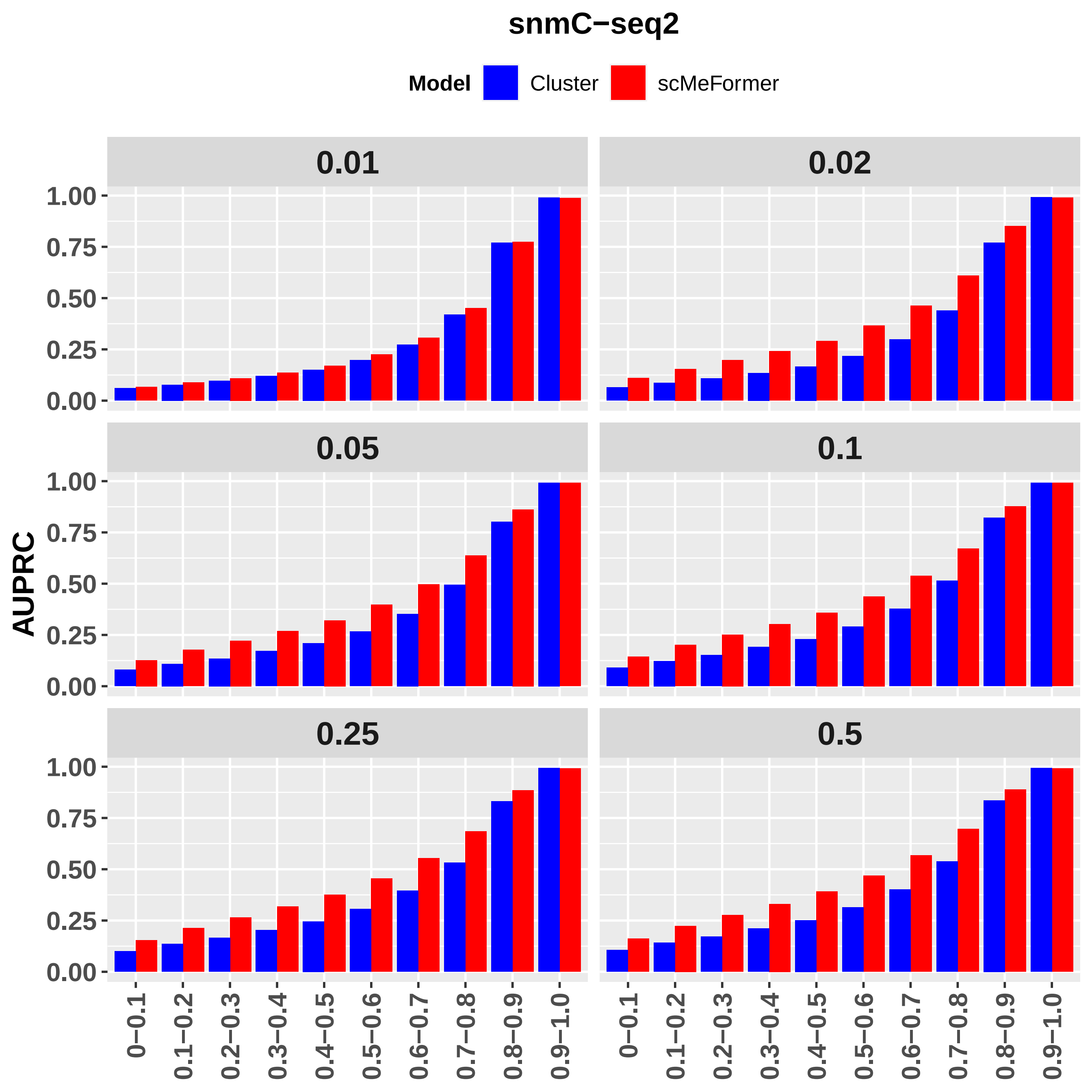

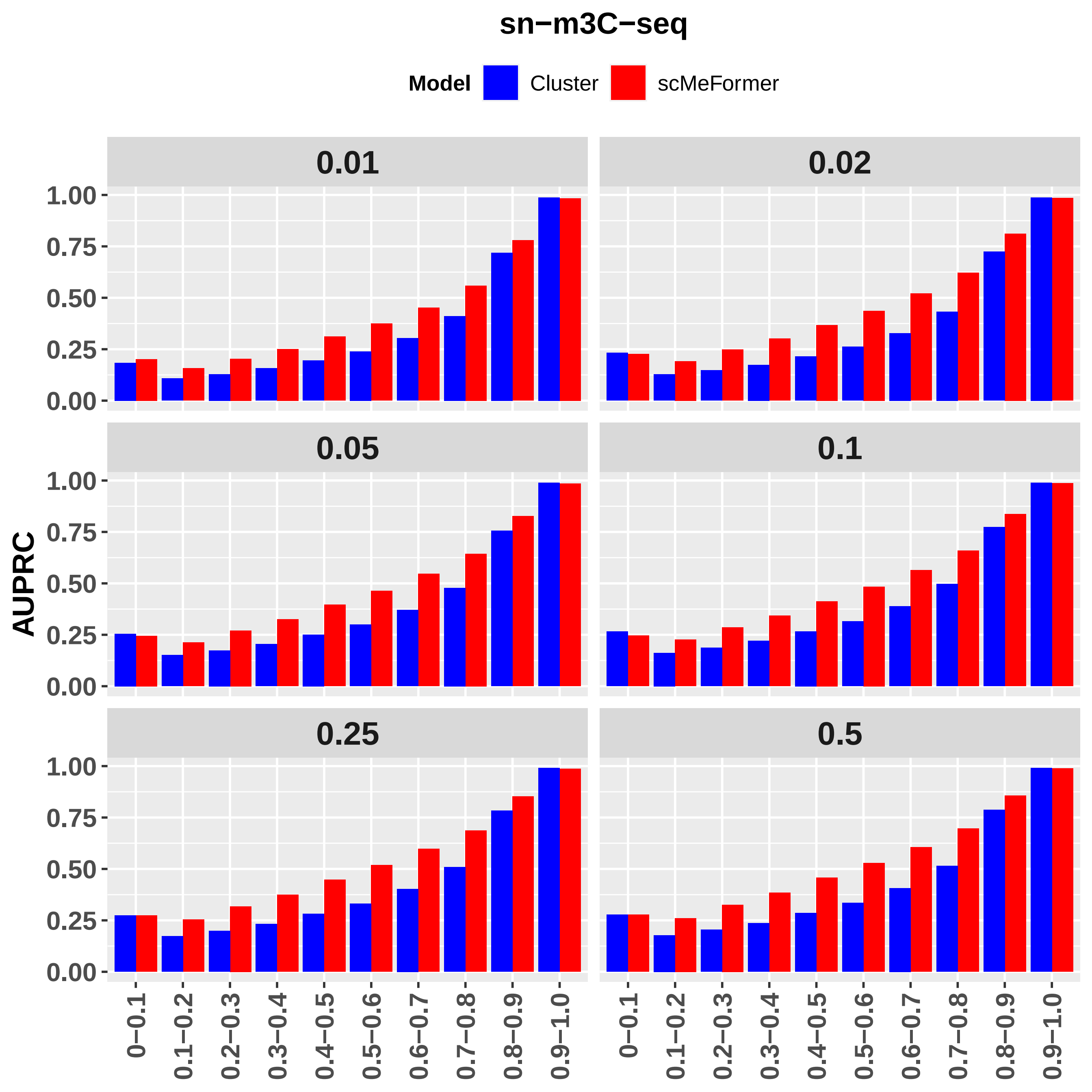

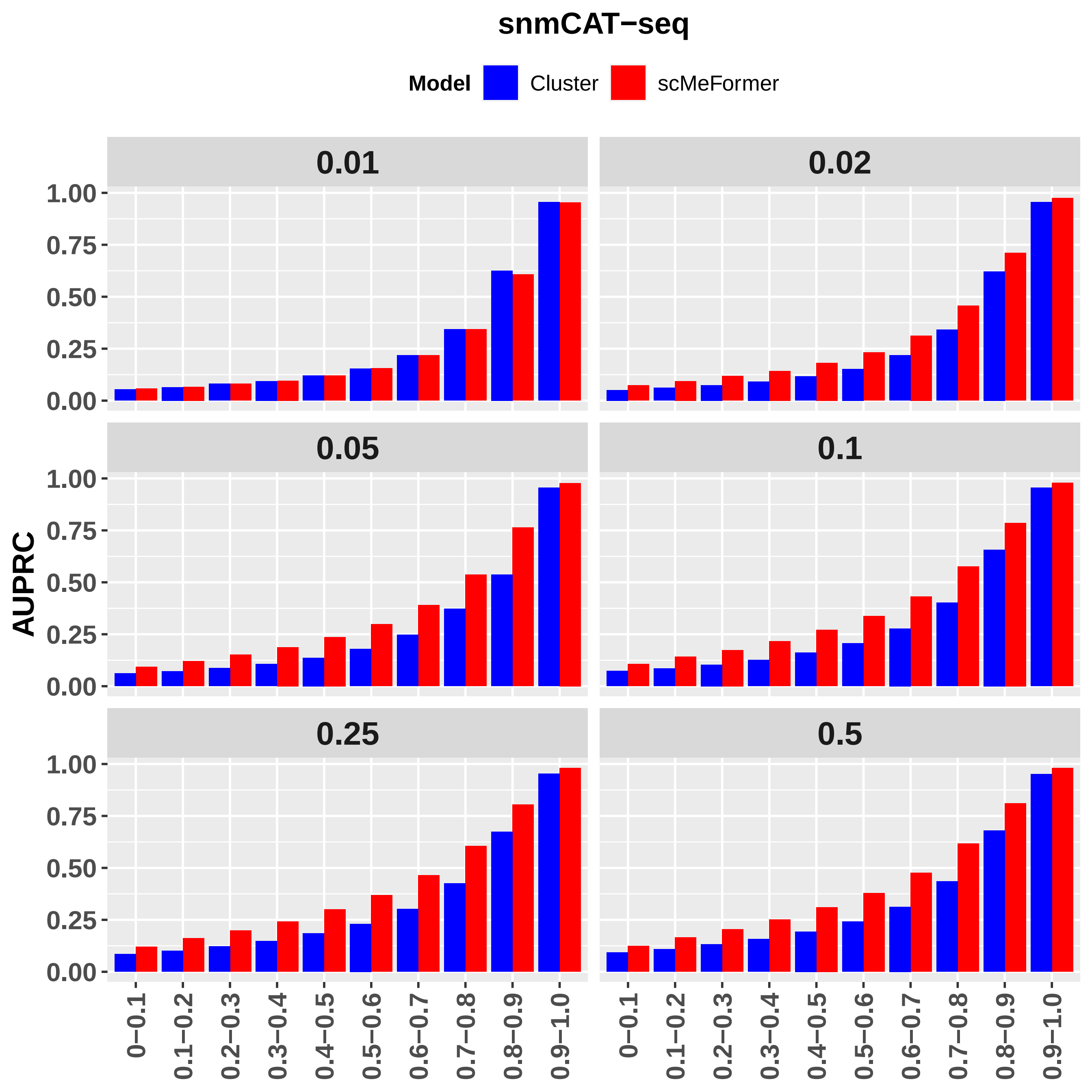

### Supplementary Figure 2

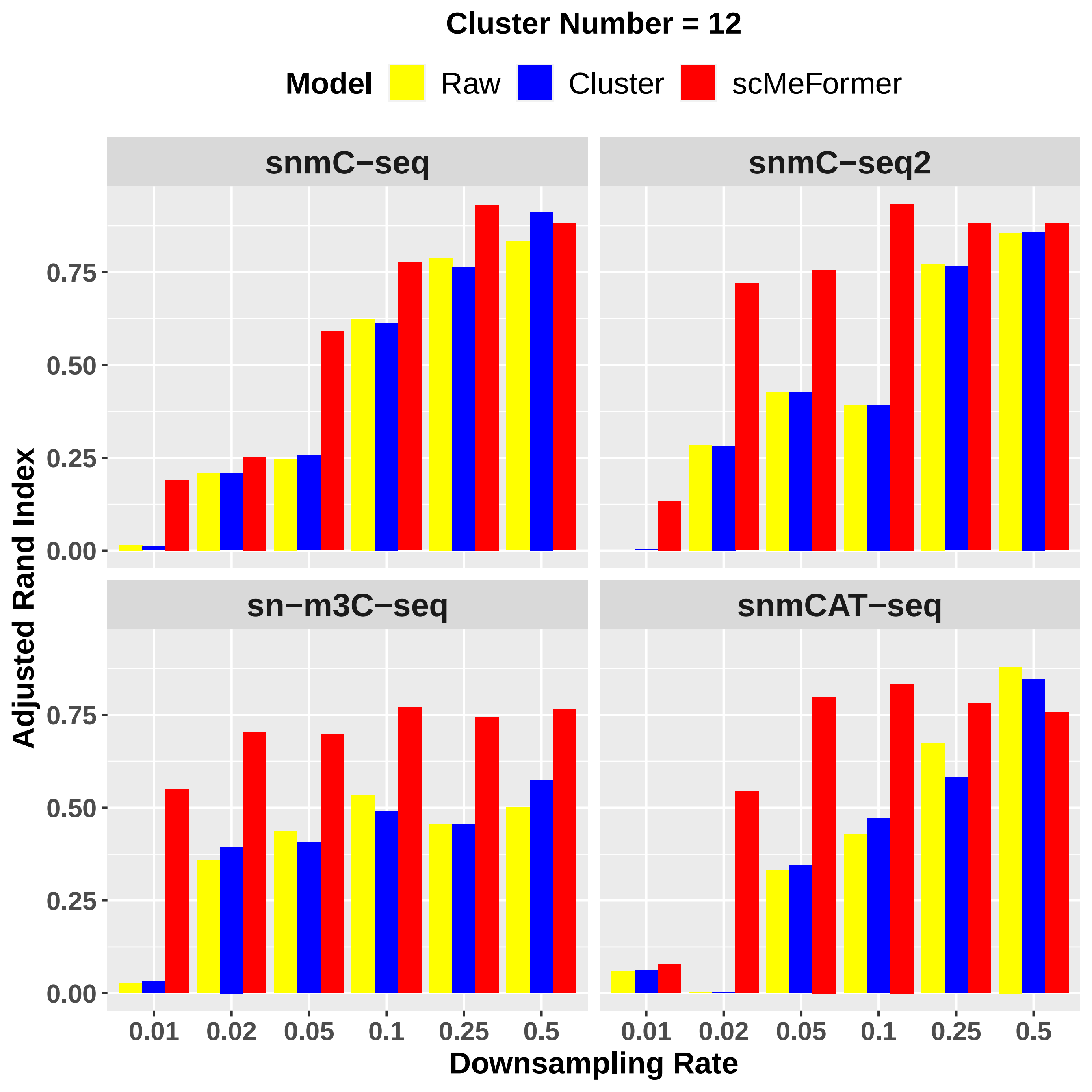

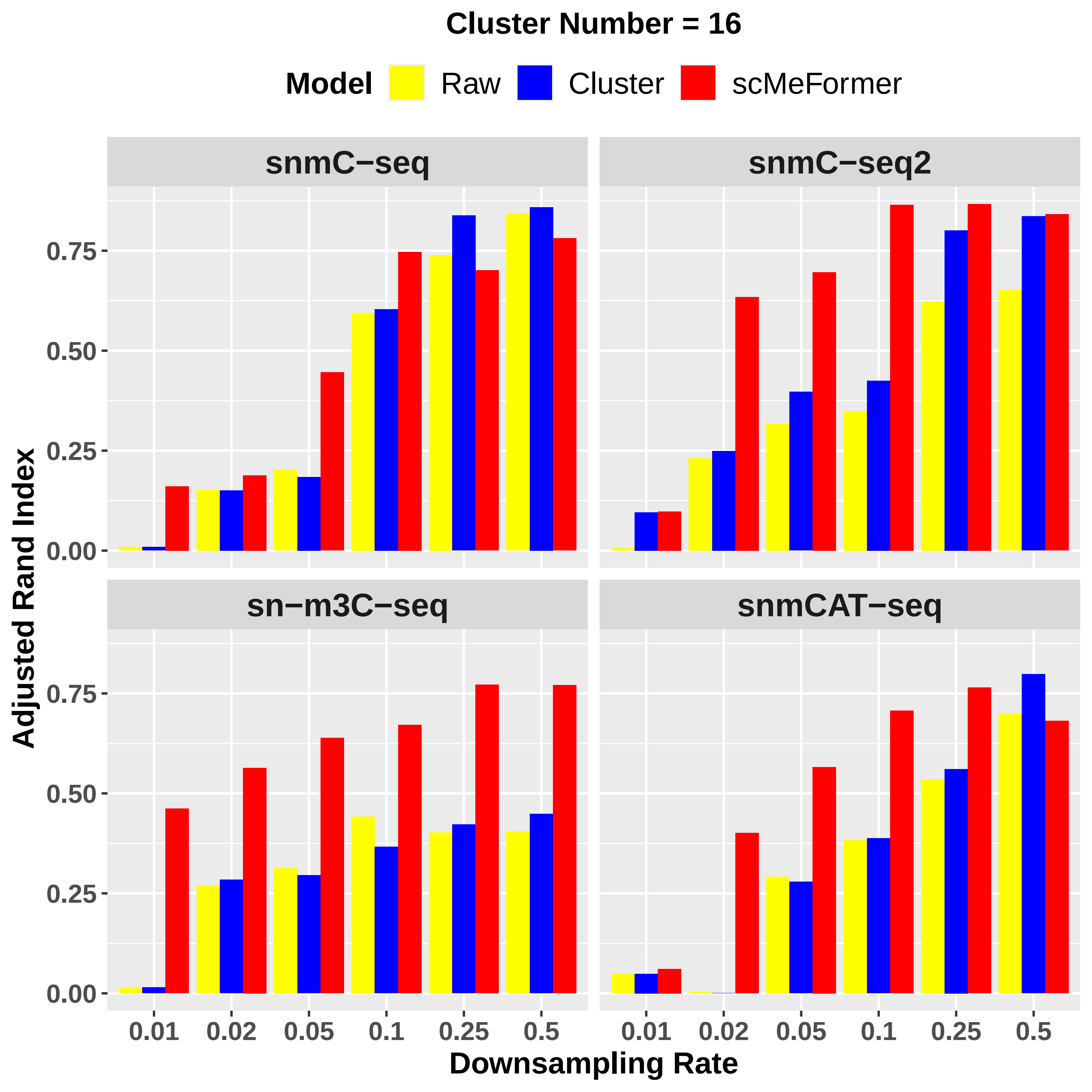

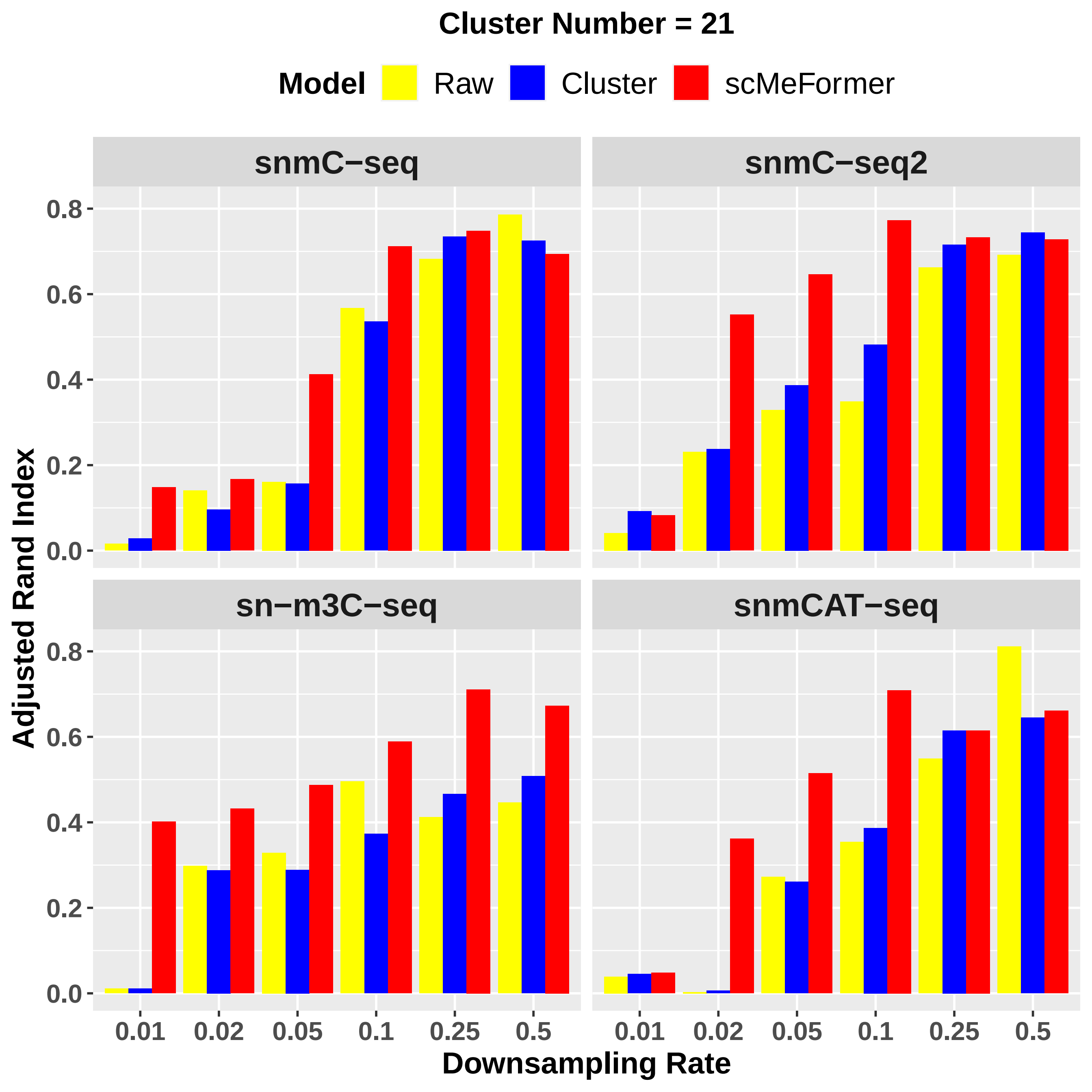

### Supplementary Figure 3

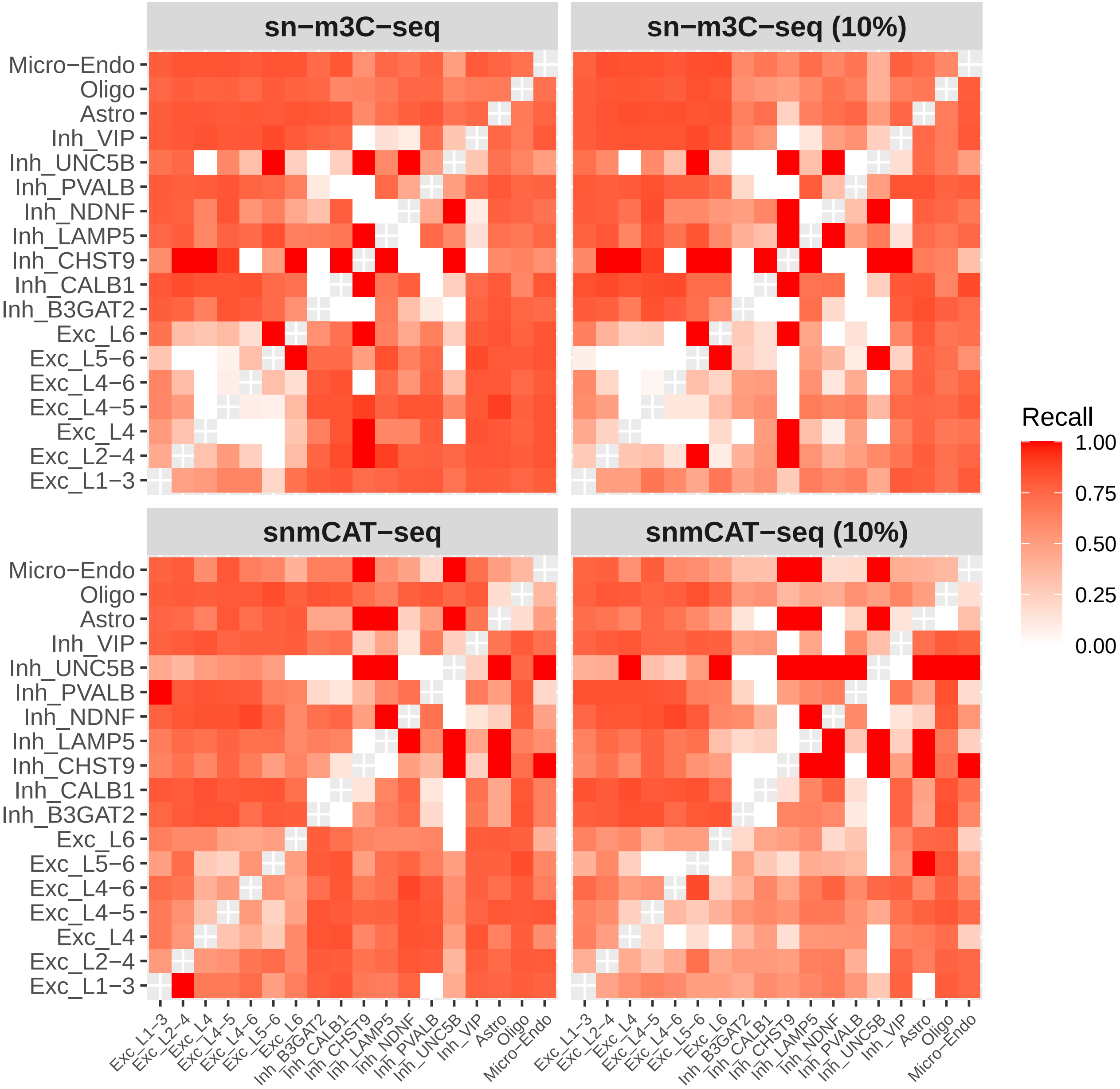

### Supplementary Figure 4

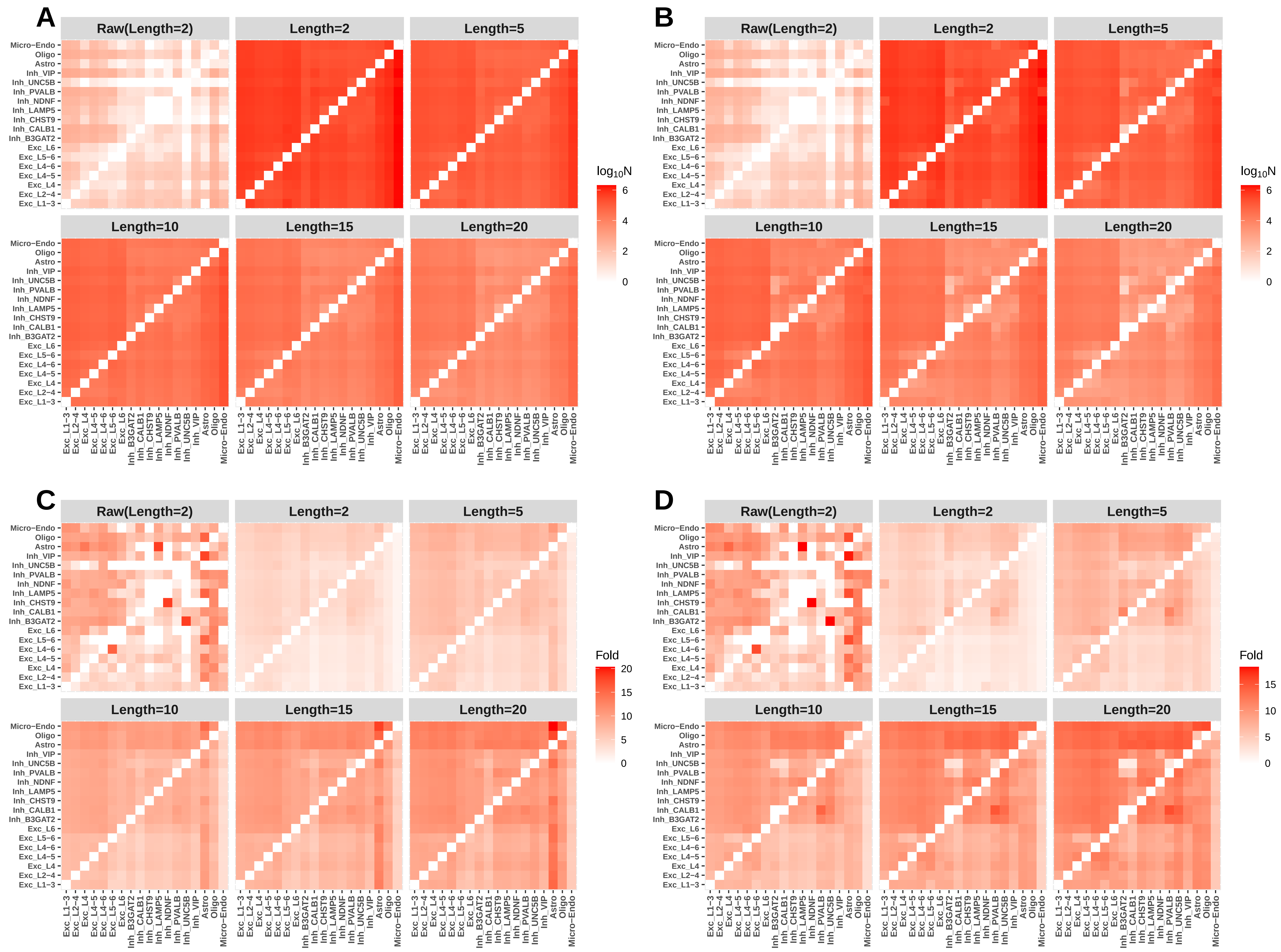

### Supplementary Figure 5

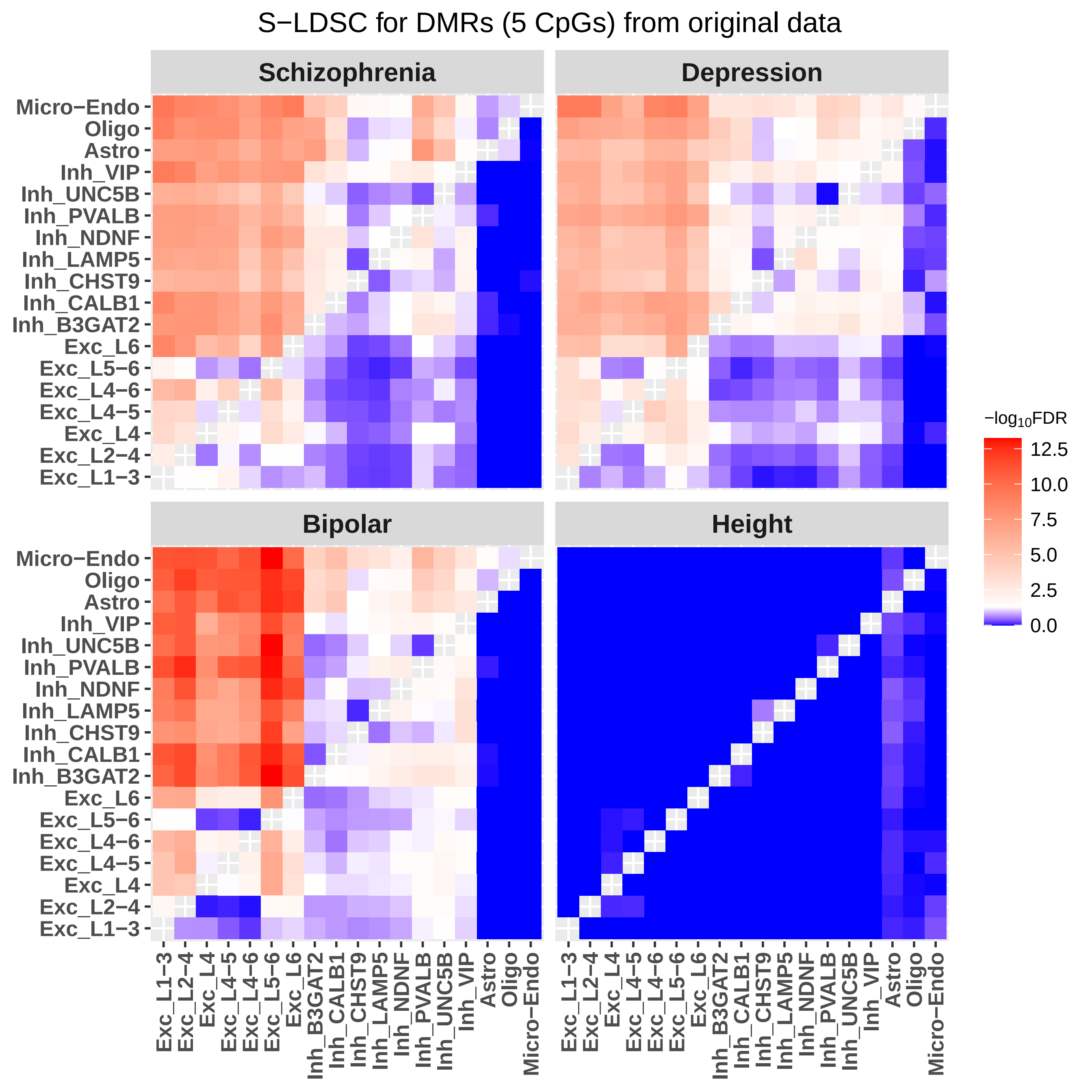

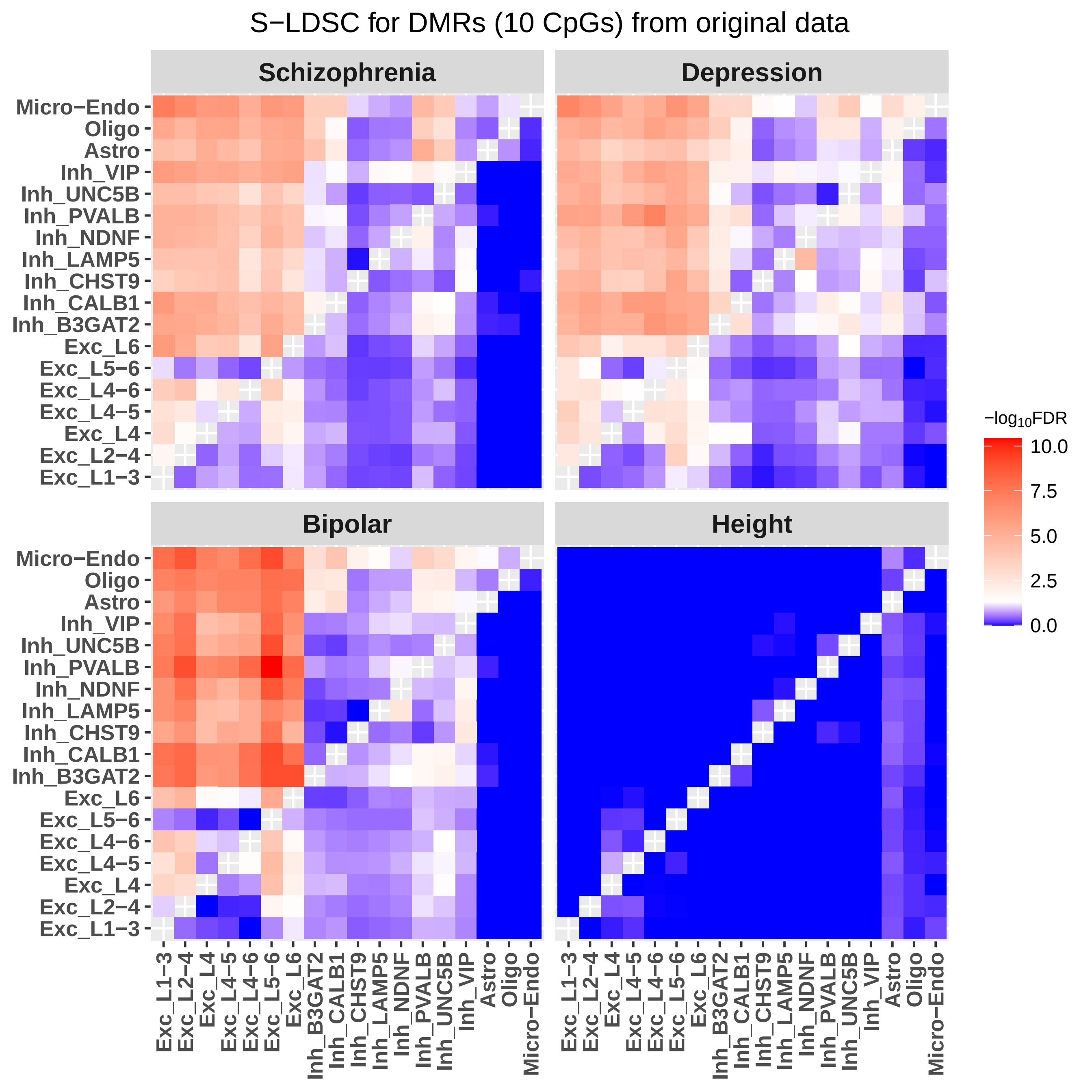

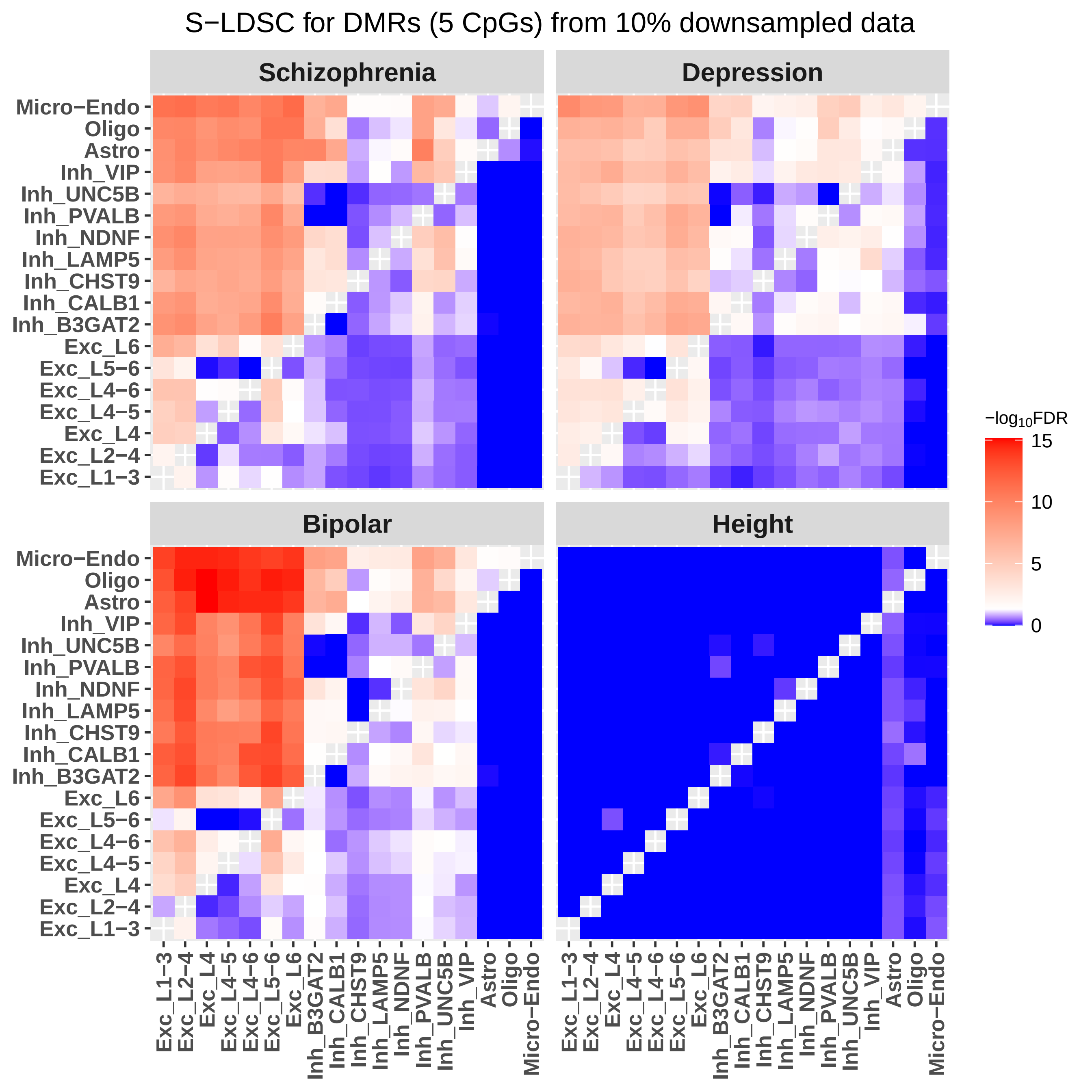

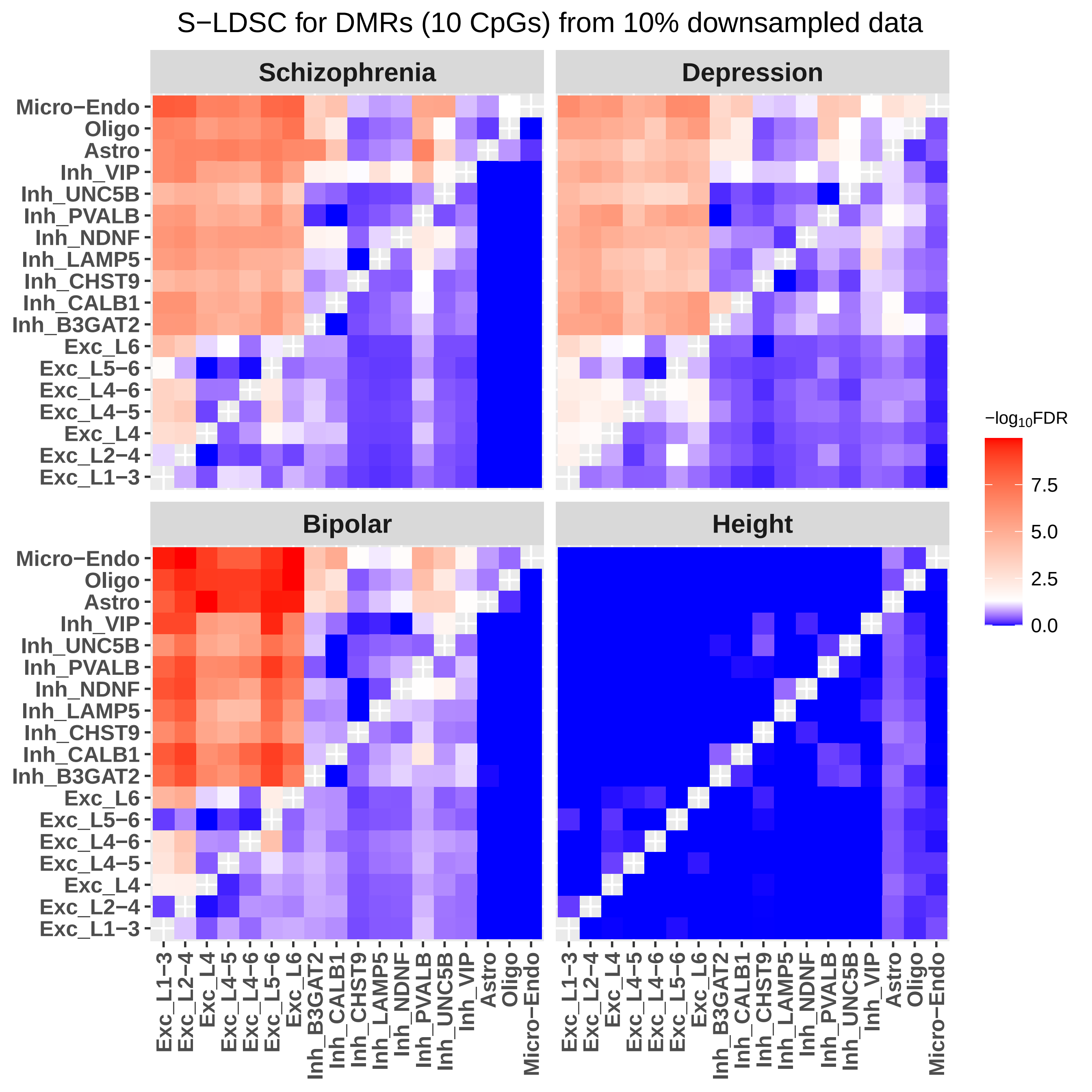

### Supplementary Figure 6

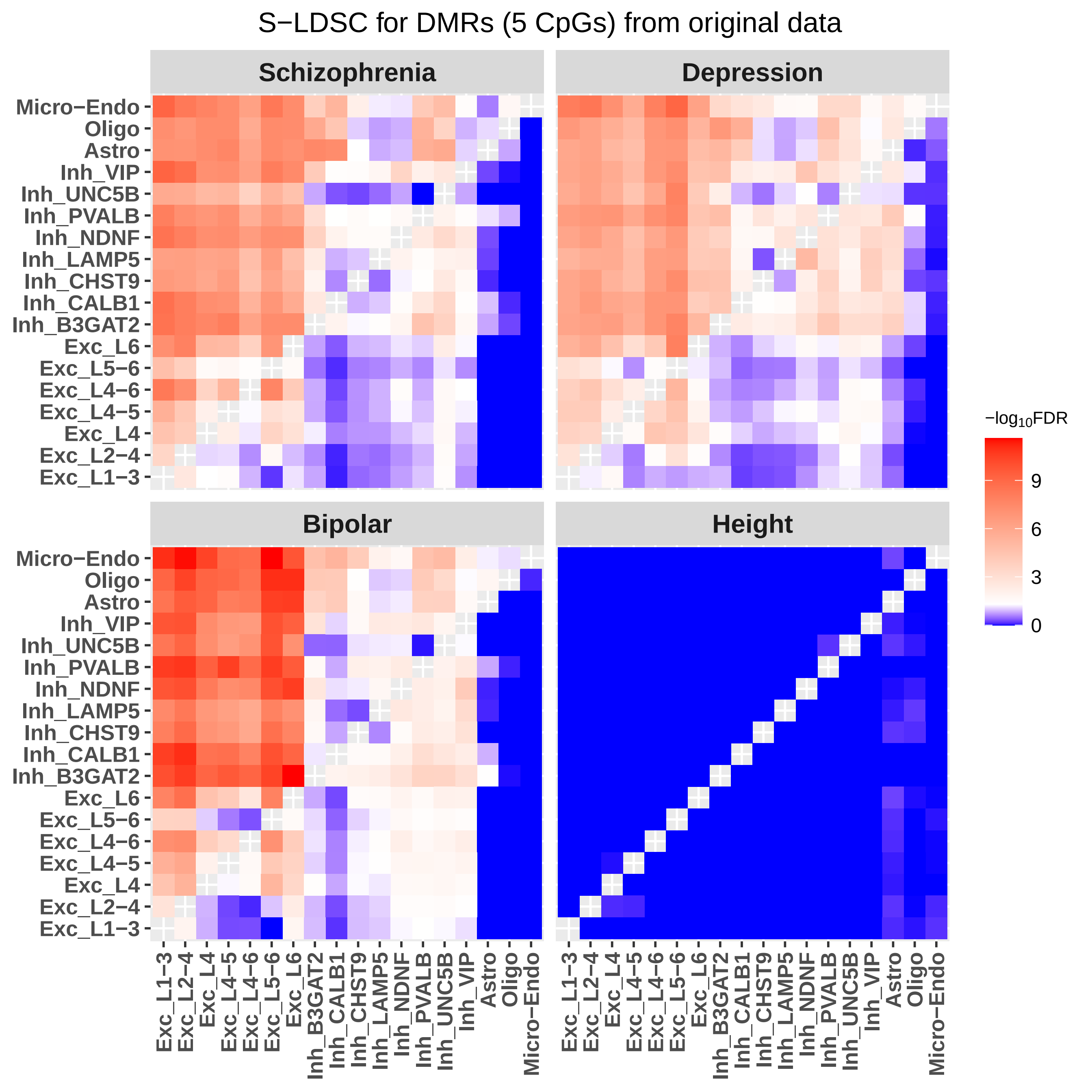

##### S−LDSC for DMRs (10 CpGs) from original data

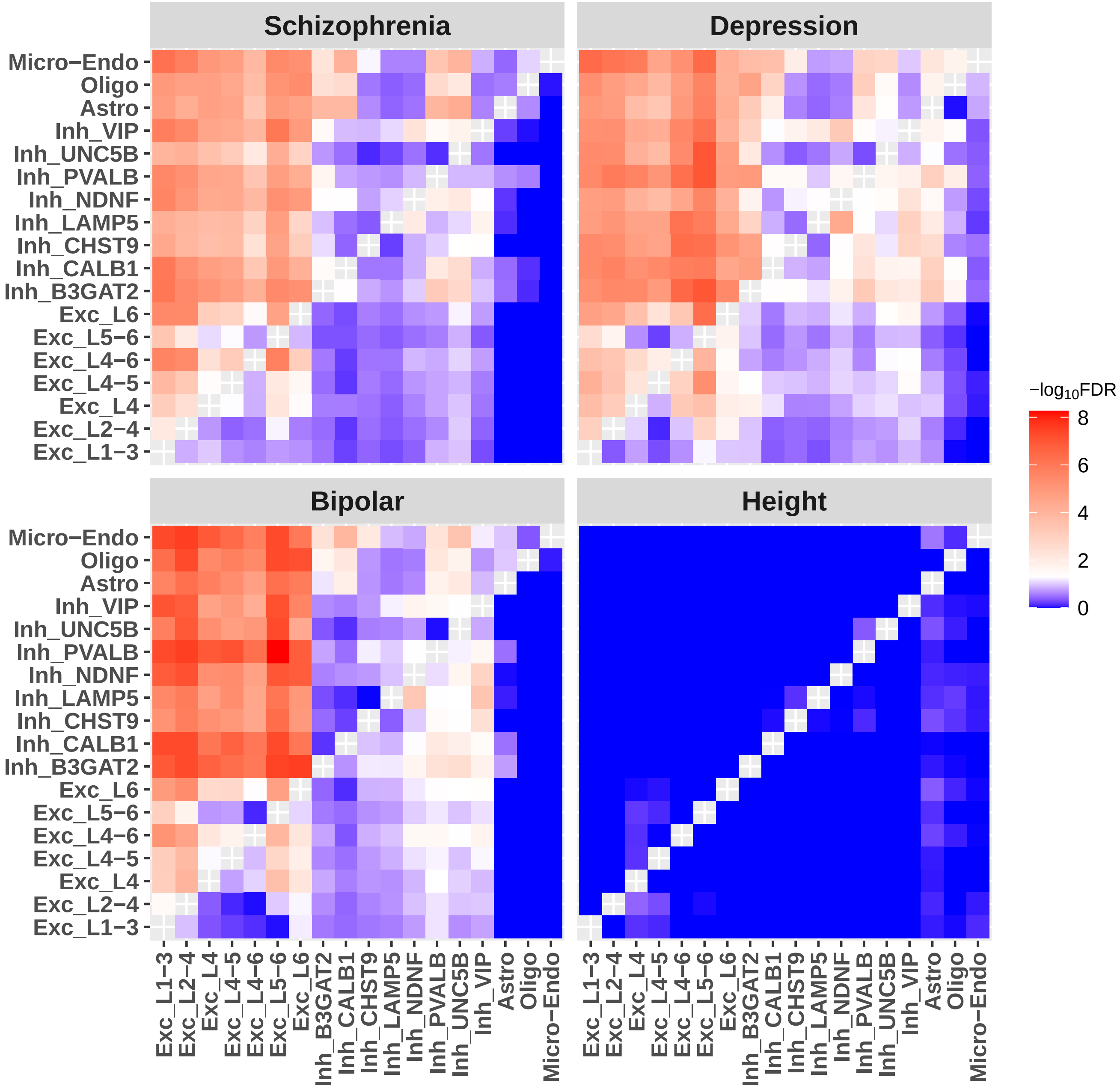

#### S−LDSC for DMRs (5 CpGs) from 10% downsampled data

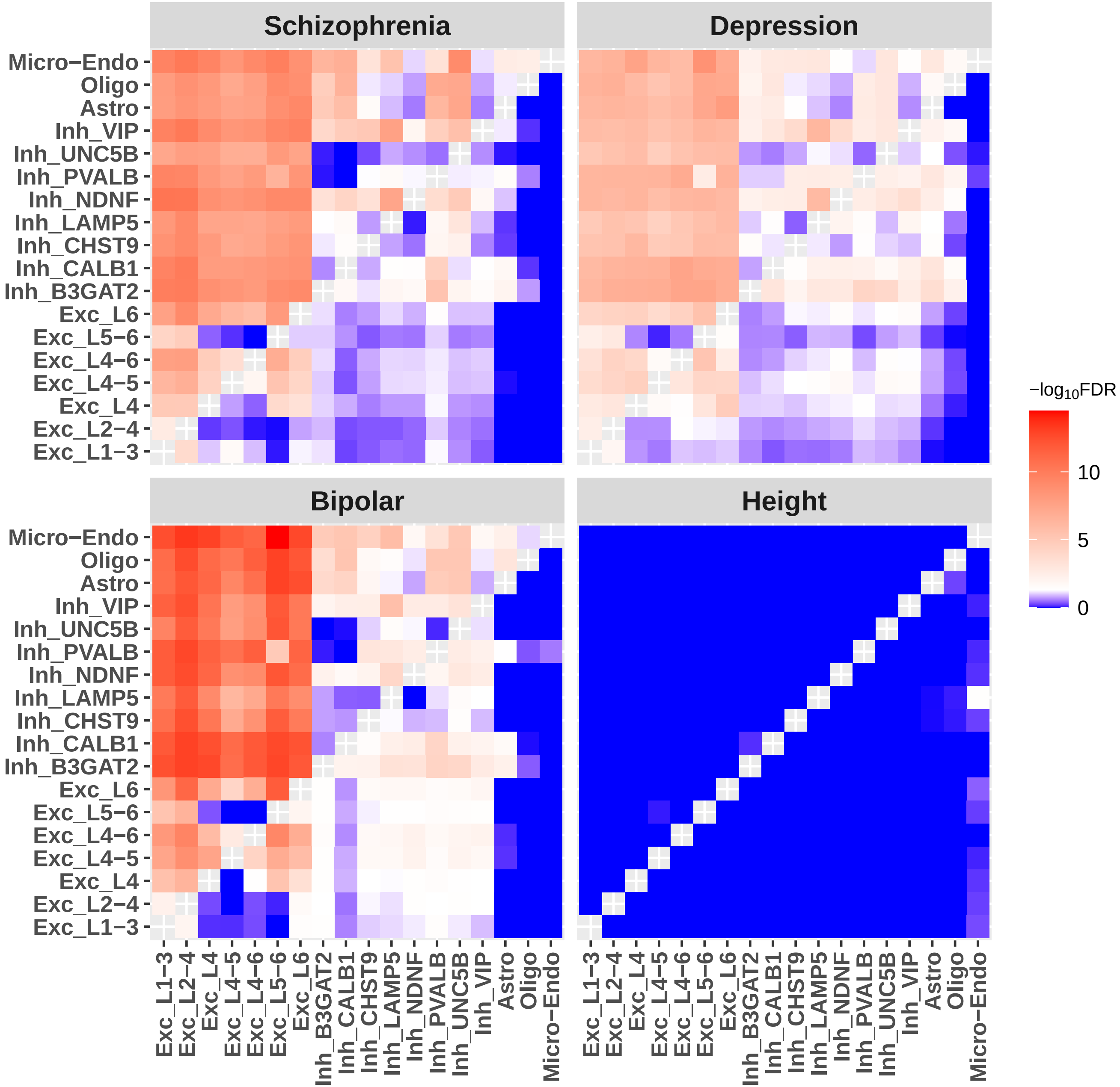

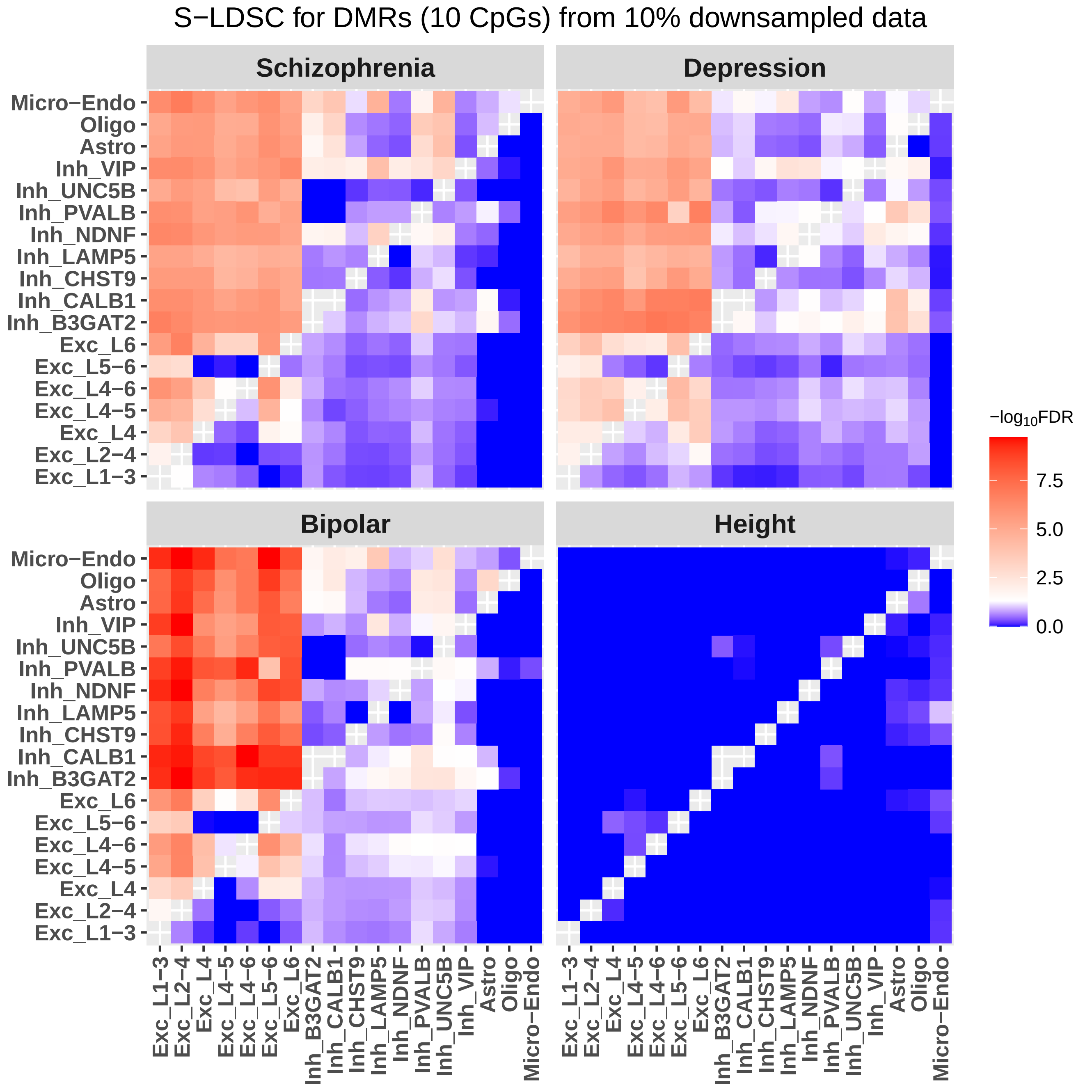

### Supplementary Figure 7

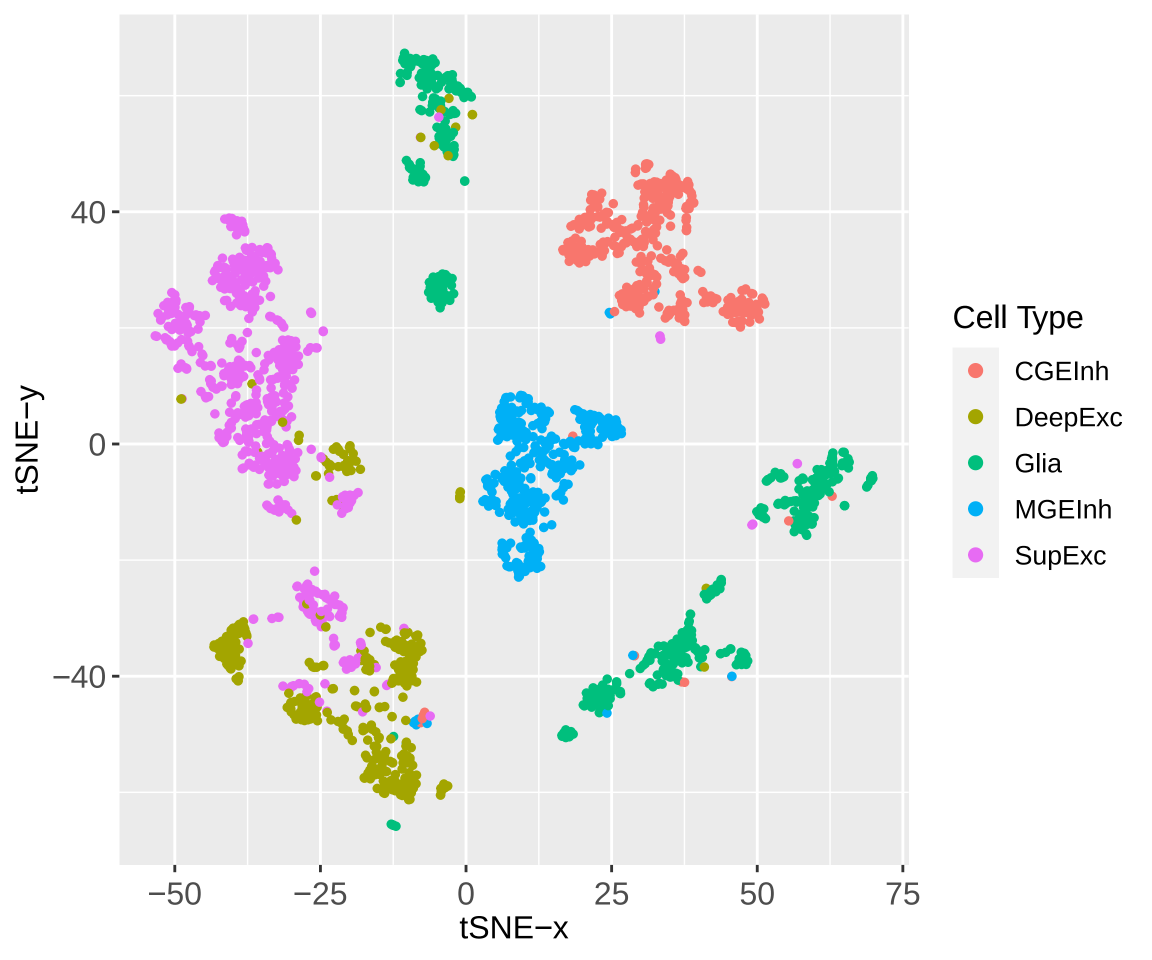
